# Profiling of microglia nodules in multiple sclerosis reveals propensity for lesion formation

**DOI:** 10.1101/2023.06.11.544204

**Authors:** Aletta M.R. van den Bosch, Marlijn van der Poel, Nina L. Fransen, Maria C.J. Vincenten, Anneleen M. Bobeldijk, Aldo Jongejan, Hendrik J. Engelenburg, Perry D. Moerland, Joost Smolders, Inge Huitinga, Jörg Hamann

**Affiliations:** Neuroimmunology Research Group, Netherlands Institute for Neuroscience, Amsterdam, The Netherlands; Department of Epidemiology and Data Science, Amsterdam Public Health Research Institute, Amsterdam University Medical Centers, Amsterdam, The Netherlands; MS Center ErasMS, Departments of Neurology and Immunology, Erasmus Medical Center, Rotterdam, The Netherlands; Swammerdam Institute for Life Sciences, University of Amsterdam, Amsterdam, The Netherlands; Amsterdam Infection & Immunity Institute, Amsterdam University Medical Centers, Amsterdam, The Netherlands

**Keywords:** multiple sclerosis, lesion formation, microglia nodules, immune activation, metabolism, mitochondria

## Abstract

Clusters of ramified HLA-DR^+^ cells, known as microglia nodules, are associated with brain pathology. Here we investigated if microglia nodules in the normal-appearing white matter (NAWM) of multiple sclerosis (MS) are different from microglia nodules in white matter (WM) in stroke and whether they may relate to the start of demyelinating MS lesions. We studied the relation between microglia nodules and pathological severity in an MS autopsy cohort (n=167), and we compared frequency, size, and gene expression of microglia nodules in MS (n=7) and stroke (n=7). MS donors with microglia nodules (64%) had a higher lesion load and a higher proportion of active lesions compared to donors without microglia nodules (36%). We found altered expression of genes in microglia nodules in MS compared to stroke, including genes previously shown to be upregulated in MS lesions. Genes associated with lipid metabolism, presence and proliferation of T and B cells, production of and response to immunoglobulins and cytokines (specifically TNF and IFN), activation of the complement cascade, and metabolic stress were upregulated. Using immunohistochemistry, we confirmed that in MS, more than in stroke, microglia nodules are associated with membrane attack complexes, have phagocytosed oxidized phospholipids, and have a tubular mitochondrial network reflecting increased metabolic activity. Furthermore, in MS, some nodules encapsulated partially demyelinated axons. Taken together, we propose that activation of some microglia nodules in MS by pro-inflammatory cytokines and immunoglobulins in combination with phagocytosis of oxidized phospholipids may lead to a volatile phenotype prone to form MS lesions.

## Introduction

Multiple sclerosis (MS) is a chronic neuroinflammatory disease characterized by focal demyelination and axonal damage throughout the brain and spinal cord ^1, 2^. Microglia are innate phagocytic glia cells of the central nervous system (CNS) that play an essential role in brain homeostasis ^3, 4^. In MS pathology, they contribute to the HLA-DR^+^ cell fraction phagocytosing myelin fragments in active and mixed active/inactive (mixed) lesions ^1, 5^. Despite many studies focusing on the role of microglia in MS, their particular role in lesion initiation is not defined yet.

In the white matter, new lesions start in the normal-appearing white matter (NAWM). Already in 1989, an magnetic resonance imaging (MRI) study showed alterations in the NAWM in MS compared to healthy control white matter (WM) ^6^, which later were in part attributed to focal microglial activation in the absence of clear demyelination ^7^. Accordingly, we recently identified subtle transcriptional changes in microglia in MS NAWM using bulk RNA sequencing ^8^. Top differentially-expressed (DE) genes related to lipid metabolism and phagocytosis that we found were also upregulated in active MS lesions, indicating early demyelination by microglia in NAWM. Since microglia adapt to local changes in the CNS ^9–11^, subpopulations with distinct cellular states may differentially contribute to MS pathology.

HLA-DR^+^ ramified microglia can accumulate and cluster in the NAWM forming small clusters of at least four up to 50 cells that are in contact with each other, which was described in relation to MS pathology for the first time in 1993 ^12, 13^. These so-called microglia nodules are regularly considered to precede MS lesion formation ^14–22^. They are found in early as well as advanced MS cases and persist throughout the disease course ^23, 24^. Moreover, they are associated with axons undergoing Wallerian degeneration ^17^ and with encapsulation of activated complement deposits ^14, 15^. Microglia nodules are engaged in phagocytosis ^25^ and express both pro-and anti-inflammatory cytokines, such as tumor necrosis factor (TNF), interleukin (IL)-1β, and IL-10 ^26, 27^. Van Horssen and colleagues reported expression of nicotinamide adenine dinucleotide phosphate (NADPH) oxidases by microglia nodules, which promotes the production of radical oxygens that can contribute to axonal damage ^27^. In sum, microglia nodules in MS express molecules involved in immune regulation and oxidative stress. However, microglia nodules are not restricted to MS, since these are also found in relation to Wallerian degeneration in brain donors with traumatic brain injury, ischemia, or stroke ^17, 22^, where microglia nodules line up around complement-opsonized axons similar as in MS ^14, 15, 22^. Therefore, to disclose MS-specific characteristics of microglial nodules and their possible contribution to MS lesion formation, we compared microglia nodules in MS with microglia nodules in stroke and with surrounding non-nodular white matter in MS and stroke. We hypothesize that in MS a subset of microglia nodules will initiate MS lesion formation. As microglia nodules in stroke are not involved in lesion formation, differences between microglia nodules in MS and in stroke may reveal mechanisms behind MS lesion formation.

We assessed the pathological and clinical relevance of presence of microglia nodules in the MS autopsy cohort of the Netherlands Brain Bank. Of microglia nodules in MS and stroke tissue, the frequency and size were quantified. Gene expression in microglia nodules and non-nodular white matter in MS and stroke was compared by RNA sequencing of laser micro-dissected tissue, and genes of interest were validated by immunohistochemistry (IHC). Using IHC, we studied the presence of lysosomal oxidized phospholipids, the presence of T and B cells, the activation of the complement cascade and formation of the membrane attack complex, the mitochondrial network in the nodules and demyelination of axons encapsulated by microglia nodules. Our gene and protein expression data show that microglia nodules in MS are different to those found in stroke. Furthermore, they indicate that part of the microglia nodules in MS show all characteristics of very small and possibly starting MS lesions. Moreover, we identify molecules and pathways that may halt the progression of microglia nodules in MS into inflammatory, demyelinating lesions.

## Materials and methods

### Characterization of MS lesions

Of 167 MS brain donors of the Netherlands Brain Bank MS cohort (NBB-MS, www.brainbank.nl), MS lesions were characterized as described previously by Luchetti and colleagues ^5^. The procedure for brain donation and the use of clinical and pathological information for research has been approved by the medical ethics committee of the VU medical center (Amsterdam, The Netherlands). Diagnoses were confirmed by a neuropathologist.

All tissue blocks (on average 23.5 ± 9.7 (standard deviation)) that were dissected during autopsy upon MRI or macroscopical appearance of lesions were stained for HLA-DR/proteolipid protein (PLP) to assess the MS lesion type ^5, 28^. The proportions of active, mixed active/inactive (mixed), inactive, and remyelinated lesions were calculated. Active lesions were defined by partial loss of PLP myelin staining and presence of HLA-DR^+^ cells throughout the lesion. Mixed lesions were defined by an inactive demyelinated center with absence of PLP staining and HLA-DR^+^ cells present at the border of the lesion. Microglia in active and mixed active/inactive lesions are stratified as ramified (score 0), rounded (score 0.5) or foamy (score 1). The microglia/macrophage activity score (MMAS) of each donor was calculated by dividing the sum of the scores of the microglia/macrophages values by the amount of active and mixed active/inactive lesions. Inactive lesions were defined by an inactive demyelinated center with no HLA-DR^+^ cells throughout. Remyelinated lesions were defined by partially myelinated axons with similar numbers of HLA-DR^+^ cells compared to the adjacent NAWM. In addition, we determined lesion load in brainstem tissue blocks as these are dissected at standard locations, which allowed us to compare the same brain region among donors ^5^. The load of reactive sites was defined as regions of accumulations of HLA-DR^+^ microglia cells in normal-appearing brainstem tissue, which are typically larger regions than nodules in which the HLA-DR^+^ microglia cells do not need to be in contact with each other to be considered as such, and total lesion load was defined as all active, mixed, inactive, and remyelinated but not reactive sites in the brainstem tissue. Additionally, each donor was scored for yes or no presence of microglia nodules in any tissue block dissected. Pathological information of MS brain donors with/without microglia nodules is summarized in **Table 1**.

**Table 1:** Donor demographics of NBB-MS cohort with and without nodules

|  | **Nodules present (all)**<br>**N=107** | **No nodules present (all)**<br>**N=60** | *p-value* |
| --- | --- | --- | --- |
| Age (years) | 63.81 (13.73) | 63.55 (12.66) | 0.90 |
| Sex | 77F, 30M | 30F, 30M | **0.01** |
| MS type | 2 RR, 30 PP, 62 SP, 13 NA | 6 RR, 20 PP, 28 SP, 6 NA | - |
| Disease duration | 28.70 (13.39) | 29.60 (12.68) | 0.83 |
| Severity score | 2.38 (0.72) | 2.37 (0.76) | 0.71 |
| Reactive site load (log) | 0.70 (0.75) | 0.32 (0.56) | **1.4e-4** |
| Lesion load (log) | 1.79 (0.97) | 1.15 (1.10) | **5.0e-4** |
| Proportion active | 0.23 (0.22) | 0.16 (0.22) | **0.03** |
| Proportion mixed | 0.30 (0.24) | 0.21 (0.25) | 0.09 |
| Proportion inactive | 0.30 (0.24) | 0.32 (0.26) | **0.03** |
| Proportion remyelinated | 0.30 (0.28) | 0.37 (0.29) | **0.02** |
| MMAS score | 0.35 (0.26) | 0.30 (0.29) | 0.15 |

Provided is the mean ± standard deviation, BRS: brainstem, F: female, M: male, NA: not available, PMD: post-mortem delay, PP: primary progressive, PR: primary relapsing, RR: relapsing-remitting, SP: secondary progressive, WM: white matter.

### Post-mortem brain tissue selection for LDM and IHC

Frozen and mirror paraffin-embedded tissue from NAWM of MS (n=7) and WM of stroke brain donors (n=7) with HLA-DR^+^ microglia nodules were matched for age, sex, post-mortem delay, and pH of the cerebrospinal fluid (CSF) (**Table 2**). Normal appearance of tissue was confirmed by intact PLP myelin staining, and microglia nodules were determined as minimally 4 HLA-DR^+^ accumulating microglia. MS brain donors that had experienced any brain infarct during life were excluded. The WM tissue of stroke brain donors did not contain infarcts. Cryo-protected frozen optic nerve samples of MS (n=4) and control donors (n=4) were previously obtained ^29^.

**Table 2:** Donor demographics of MS and stroke donors

|  | **Age**<br>**(years)** | **Sex** | **PMD**<br>**(h:min)** | **pH of CSF** | **MS type** | **Disease**<br>**duration**<br>**(years)** | **Time to**<br>**EDSS6**<br>**(years)** | **Time of**<br>**stroke to**<br>**death**<br>**(months)** |
| --- | --- | --- | --- | --- | --- | --- | --- | --- |
| MS (*n*=7) | 66 (12) | 5M/2F | 8:50 (7:19) | 6.47 (0.10) | 6 SP/1 PP | 38 (12) | 26 (15) | 64 (111) |
| Stroke (*n*=7) | 80 (10) | 4M/3F | 7:18 (4:38) | 6.42 (0.24) | - | - | - | - |
| *p-value* | 0.06 | 0.58 | 0.43 | 0.59 |  |  |  |  |

PMD: post-mortem delay, CSF: cerebrospinal fluid. Data presented as average with standard deviation.

### Immunohistochemistry

For IHC, (NA)WM tissue of MS (n=9) and stroke (n=8) brain donors was cut from frozen (20 µm) or paraffin-embedded (8 µm) tissue blocks not containing any lesions or reactive sites. For SMI312 and PLP, normal-appearing optic nerve tissue of MS donors was fixed overnight in 4% paraformaldehyde, protected in 30% sucrose for 24 hours, frozen, and cut at 20 µm ^29^. For paraffin-embedded tissue, antigen retrieval was performed as indicated in **Table 3** for 10 min in a microwave at 700 W. Sections were incubated overnight at 4°C with primary antibodies (**Table 3**) diluted in incubation buffer (for paraffin tissue: 0.5% gelatin and 0.5% Triton X-100 in TBS; for frozen tissue: 1% bovine serum albumin and 0.5% Triton X-100 in phosphate-buffered saline, pH7.6). HLA-DR-and PLP-stained sections were incubated with HRP-labelled anti-mouse antibody (K5007, Dako Real EnVision detection system; Dako, Nowy Sącz, Poland) for 1 hour and immunostaining was visualized with 3’3-diaminobenzidine (1:100; Dako) incubation for 10 min, followed by counterstaining in haematoxylin for 30 sec and mounted in Entellan (Merck, Kenilworth, NJ, USA). For CD38-stained sections, CD38 was incubated with avidin-biotin complex – alkaline phosphatase kit (Vector, Olean, New York, United States) (1:800) for 1 hour and staining was visualized with the ImmPACT Vector Red Alkaline Phosphatase Substrate Kit (Vector). For fluorescent stainings, the sections were either incubated with a compatible fluorophore (1:400), or the staining was enhanced with a compatible biotinylated secondary antibody (1:400) for 1 hour followed by avidin-biotin complex – HRP kit (1:800) for 45 min, biotinylated tyramide (1:10,000) for 10 min, ABC (1:800) for 45 min at RT, and lastly, streptavidin-conjugated fluorophore (1:800) for 1 hour. All fluorescent stainings were incubated with Hoechst (1:1,000) for 10 min, 0.1% Sudan Black in 70% ethanol for 10 min and mounted in Mowiol.

**Table 3:** Antibodies overview

| **Antigen** | **Supplier (cat#)** | **Clone** | **Dilution** | **Antigen retrieval** |
| --- | --- | --- | --- | --- |
| C1qB | Abcam (ab92508) | EPR2981 | 1:100* | Tris EDTA buffer pH9 |
| C3d | Dako (A0063) | Polyclonal | 1:300 | Citrate buffer pH6 |
| C59b | Dako (M077701-8) | aE11 | 1:100 | Citrate buffer pH6 |
| CD138 | BioRad (MCA2459T) | B-A38 | 1:250 | Citrate buffer pH6 |
| CD20 | Dako (M0755) | L26 | 1:100 | Citrate buffer pH6 |
| CD3 | Dako (A0452) | Polyclonal | 1:100 | Citrate buffer pH6 |
| CD38 | Atlas antibodies (HPA022132) | Polyclonal | 1:3000 | Citrate buffer pH6 |
| CD4 | Dako (M7310) | 4B12 | 1:100 | Tris EDTA buffer pH9 |
| CD8 | BD Biosciences (641400) | SK1 | 1:500 | Citrate buffer pH6 |
| DAGLB | Atlas Prestige (HPA069377) | Polyclonal | 1:50* | PBS pH7.6 |
| E06 | Avanti (330002S) | T15 | 1:100 | PBS pH7.6 |
| FABP5 | RabMab (ab255276) | EPR22552-64 | 1:2000 | Tris EDTA buffer pH9 |
| HLA-DR | Dako (M0775) | CR3 | 1:100 | Citrate buffer pH6 |
| IAH1 | Invitrogen (PA5-65270) | Polyclonal | 1:50* | PBS pH7.6 |
| Iba1 | Wako (019-19741) | Polyclonal | 1:500 | Citrate buffer pH6 |
| LAMP1 | Abcam (ab24170) | Polyclonal | 1:200 | Citrate buffer pH6 |
| MBP | Sigma (AB980) | Polyclonal | 1:200 | Citrate buffer pH6 |
| PCNA | Santa Cruz (sc-25280) | PC10 | 1:1000 | Citrate buffer pH6 |
| PLP | Secotec (MCA839G) | plpc1 | 1:3000 | Citrate buffer pH6 |
| SMI312 | Eurogentec (SMI-312R) | SMI-312R | 1:6000 | Citrate buffer pH6 |
| STARD13 | Invitrogen (PA5-63622) | Polyclonal | 1:300* | PBS pH7.6 |
| Tomm20 | RabMab (ab186735) | EPR15581-54 | 1:100* | Citrate buffer pH6 |

\*Staining enhanced

### Quantification of immunohistochemistry

HLA-DR^+^ microglia nodules were visualized with an Axioplan2 microscope (Zeiss, Oberkochen, Germany). The entire tissue section was scanned to manually count nodule numbers in each section and corrected for size of the tissue section. To determine the size of microglia nodules, a picture was made of each nodule and a macro to automatically determine nodule size was developed using Image-Pro software (MediaCybernetics, Bethesda, MD, USA). An outline of each nodule was drawn manually, and an area mask was placed to capture HLA-DR-stained microglia nodules using a greyscale intensity threshold >50 for HLA-DR/PLP stainings and a threshold >110 for HLA-DR stainings. The total area of HLA-DR-stained microglia nodules was automatically calculated and expressed as µm^2^.

For CD138, CD3, CD20, CD4, CD8, and PCNA, stainings were visualized using a confocal laser-scanning microscope (SP8; Leica, Wetzlar, Germany) with the software LASX, magnification 40x. Each tissue section was scanned for HLA-DR^+^ or IBA1^+^ microglia nodules, and pictures were made with each nodule present in the middle to detect immune cells around microglia nodules in a radius of 150-180 µm. Pictures were processed and analyzed using Fiji software ^30^. For CD38, C1qB, C3d, MAC, FABP5, DAGLB, IAH1, and STARD13, scans were made on the Axio slide scanner, x20 magnification (ZEISS, Oberkochen, Germany). Of each section, all HLA-DR^+^ microglia nodules were annotated as positive or negative for C1qBb, MAC, FABP5, DAGLB, IAH1, and STARD13, and for CD38 nodules were annotated as positive or negative for CD38^+^ cells in a radius of 150-180 µm on Qupath (version 0.4.3). The SMI312, PLP, and HLA triple staining, the LAMP1, E06, and HLA triple staining and the Tomm20 and HLA double staining were visualized at 63x using stimulated emission depletion (STED) microscopy (STEDYCON; Abberior Instruments, Göttingen, Germany) for z-stacks. 3D-rendered images were analyzed for (partial) demyelination in Fiji. For PLP triple staining, E06 triple staining and Tomm20 double staining, z-stack images were taken of all HLA^+^ microglia nodules in each section on the STED microscope at magnification 63x with 0.5-µm step size. For E06, using Fiji plugin for ImageJ, all microglia nodules were annotated as positive or negative for LAMP1^+^ E06^+^ phagocytosed oxidized phospholipids within the nodule. For Tomm20, the length of the mitochondria network was measured for each nodule using Fiji. The mitochondrial network inside the nodule was considered as fragmented if all mitochondria were <2 µm, intermediate if the largest mitochondria was between 2-5 µm, and tubular if the largest mitochondria was >5 µm in size ^31^.

### Laser capture microscopy and RNA isolation

Frozen tissue sections (20 µm) were mounted on PARM MembraneSlides (P.A.L.M. Microlaser Technologies, Bernried, Germany) and dried for 48 hours at room temperature in a sealed box containing silica gel. Sections were fixed in ice-cold dehydrated acetone on ice for 10 min and dried at RT in a slide box with silica gel. Sections were incubated with biotinylated HLA (1:100) and RNase inhibitor (1:500) in PBS + 0.5% Triton X-100 for 15 min and ABC (1:800) with RNase inhibitor (1:500) in PBS for 15 min. Immunostaining was visualized with 3’3-diaminobenzidine (1:100; Dako) incubation for 5 min at RT, followed by counterstaining in cresylviolet (0.1% in 70% EtOH) for 10 seconds. A series of 70%–86%–100% ethanol was dripped over the section and sections were transferred to the laser dissection microscope (LDM) (ZEISS) immediately.

Of the selected donors, 90-151 microglia nodules and an equal amount of non-nodular (NA)WM tissue were collected from 8-22 sections, depending on the nodule frequency. Tissue was lysed in 50 µl Trisure (Bioline, London, UK), and RNA was isolated with the RNeasy Micro Kit (Qiagen) using an adapted protocol. Chloroform was added 1:5, and samples were vortexed and incubated on ice for 5 min. After centrifugation at 11,000 rpm for 15 min at 4˚C, the aqueous phase was transferred to a new tube. The remaining sample was incubated with 1 volume of chloroform, vortexed, and centrifuged at 11,000 rpm for 15 min at 4˚C. The aqueous phases were combined. 1 volume of 70% EtOH was added and the sample was transferred to the loading column. The sample was run through the column 3 times by centrifugation for 20 sec at 10,000 rpm at 4˚C. RW1 buffer was added to the column and centrifuged at 10,000 rpm for 20 sec at 4˚C. DNAse1 and RDD buffer were incubated for 15 min RT. The sample was washed with RPE buffer followed by 70% EtOH for 20 sec at 10,000 rpm, and the column was allowed to dry for 5 min at 14,000 rpm. 14 µl of RNase-free H2O was added to the column, incubated at RT for 2 min, and RNA was collected by centrifugation for 1 min at 14,000 rpm.

### RNA sequencing and gene expression analysis

PolyA-enriched mRNA sequencing on a Illumina NovaSeq6000 system and sequence alignment were performed by GenomeScan (Leiden, The Netherlands). Putative adapter sequences were removed from the reads when the bases matched a sequence in the TruSeq adapter sequence set using cutadapt (v2.10). Trimmed reads were mapped to the human reference genome GRCh37.75 using HiSAT2 v-2-1.0 ^32^). Gene level counts were obtained using HTSeq (v0.11.0) ^33^. Statistical analyses were performed using the edgeR ^34^ and limma/voom ^35^ R/Bioconductor packages (R: v4.0.0; Bioconductor: v3.11). Seven highly abundant mitochondrial genes were removed from the dataset. Genes with more than 2 reads in at least 4 of the samples were retained. Count data were transformed to log2-counts per million (logCPM), normalized by applying the trimmed mean of M-values method ^36^, and precision weighted using voom ^37^. One stroke donor (nodule and non-nodular WM sample) and one MS non-nodular NAWM sample were identified as outliers and removed from the dataset **(Supplementary Table 1)**. Differential expression was assessed using an empirical Bayes moderated t-test within limma’s linear model framework including the precision weights estimated by voom and the consensus correlation between samples of the same donor (function ‘duplicateCorrelation’, limma package). The differential expression analysis was performed both with and without a covariate for the estimated microglia content (in percent). The proportion of microglia was determined using cell type deconvolution with dtangle ^38^ using the set of markers from Darmanis et al. ^39^ and using the script DeconvAnalysis.Rmd by Patrick et al. ^40^ as a template. To test for the possible presence of lymphocytes, we also performed cell type deconvolution using lymphocyte markers as reported by Schirmer et al. ^41^ and Palmer et al. ^42^. Resulting p values were corrected for multiple testing using the Benjamini-Hochberg false discovery rate. Genes were re-annotated using biomaRt using the Ensembl genome databases (v103). All differentially expressed genes with an adjusted p value of <0.05 were sorted on logFC, and up to 50 top differentially expressed genes were summarized with the p value and logFC adjusted for microglia proportion indicated. Principal component analysis (PCA) was performed on the logCPM values of the 500 most variable genes to distinguish sources of variation. Functional annotation and gene ontology (GO) analysis was performed on genes significantly differentially expressed with adjusted p<0.05 between groups using DAVID ^43^, with Homo sapiens as background set. As GO terms were utilized to find genes of interest, no multiple testing correction was performed.

### qPCR of genes involved in immunoglobulin production

Frozen tissue section (20 µm) was cut from stroke (n=6) and MS (n=6) (NA)WM tissue containing microglia nodules and lysed in 800 µl TRIsure. RNA was isolated according to manufacturer’s instructions (Bioline). Briefly, chloroform (1:5) was added to each TRIsure sample and after centrifugation, the aqueous phase was collected, followed by incubation with 1 µg glycogen (Roche, Basel, Switzerland) for 30 min in ice-cold isopropanol at −20˚C. Precipitated RNA was washed in ice-cold 75% ethanol and diluted in 20 µl deionized water.

Synthesis of cDNA was performed according to manufacturer’s instructions, using the Quantitect Reverse Transcription kit (Qiagen, Hilden, Germany). 50 ng RNA was mixed with 1 µl gDNA Wipe-out buffer and incubated for 2 min at 42°C, followed by incubation with QuantiTect Buffer, RT Primer Mix, and Quantitect Reverse Transcriptase for 30 min at 42°C and incubation for 3 min at 95°C.

To determine gene expression of immunoglobulin genes, quantitative polymerase chain reaction (qPCR) was performed. Control WM tissue and tissue collected from MS lesions and lymph node was used as negative and positive control samples, respectively. Primers were designed at the Integrated DNA Technologies website (eu.ifdna.com). Optimal primers were selected based on dissociation curve and specificity, examined on cDNA derived from control, MS NAWM, and MS lesioned tissue. Gene expression was normalized on the mean of two housekeeping genes, *GAPDH* and *EEF1A1*. For each gene, the relative expression was calculated using the 2^-ΔΔCT^ method. Primers used for reverse transcription (RT)-qPCR are provided in **Supplementary Table 2**.

### Statistical analysis

Data obtained from immunohistochemistry and RT-qPCR was tested for normality by a Shapiro-Wilk normality test, followed by parametric or non-parametric tests for numeric data, quasibinomial generalized linear mixed models for proportional data, or a chi square test for binomial data to define p-values. Statistical analyses were performed in Rstudio (version 1.2.5033; Rstudio, Boston, MA, USA) for R (version 4.2.0), using key packages ggplot2, lme4, car, plyr, ggpubr, Hmisc, and corrplot. P values <0.05 were considered significant.

## Results

### Microglia nodules in MS correlate with MS pathology

In the MS cohort of the NBB, we quantified the number of MS donors with and without microglia nodules. To determine the relevance of microglia nodules for MS pathology and clinical course, we compared the pathological and clinical characteristics of MS donors with and without microglia nodule in all blocks dissected. Out of 167 MS brain donors, 107 donors (64%) had microglia nodules present in at least one tissue block dissected, and 60 donors (36%) did not have any microglia nodules in any of the tissue blocks dissected. MS donors with microglia nodules present in at least one tissue block, compared to MS donors without microglia nodules, had a significantly higher number of reactive sites (log, with nodules: 0.70 ± 0.75, without nodules: 0.32 ± 0.56, p=6.0e-4) and higher lesion load (log, with nodules: 1.79 ± 0.97, without nodules: 1.15 ± 1.10, p=4.5e-4) in standard locations, a higher proportion of active lesions (with nodules: 0.23 ± 0.22, without nodules: 0.16 ± 0.22, p=0.03) and a lower proportion of inactive lesions (with nodules: 0.30 ± 0.24, without nodules: 0.32 ± 0.26, p=0.03) and remyelinated lesions (with nodules: 0.30 ± 0.28, without nodules: 0.37 ± 0.29, p=0.02) compared to MS donors without microglia nodules present. There was no difference in proportion of mixed active/inactive lesions, MMAS, disease severity measured as time to expanded disease disability scale (EDSS) 6, or disease duration (**Table 1**).

### Microglia nodules in MS are more frequent than in stroke but are not different in size

Nodule frequency and size were quantified in (NA)WM tissue sections of MS (n=7) and stroke donors (n=7). We observed a significantly higher number of microglia nodules in MS NAWM tissue as compared to stroke WM (MS: 18.1 ± 15.5 per 100 mm^2^, stroke: 4.8 ± 2.2 per 100 mm^2^, p=0.03), but the microglia nodules were not different in size (p=0.16, **Fig. 1a-c**). Furthermore, in contrast to stroke WM tissue with microglia nodules, the majority of MS NAWM tissue with microglia nodules showed many HLA-DR^+^ ramified microglia throughout the whole tissue section.

**Figure 1:**
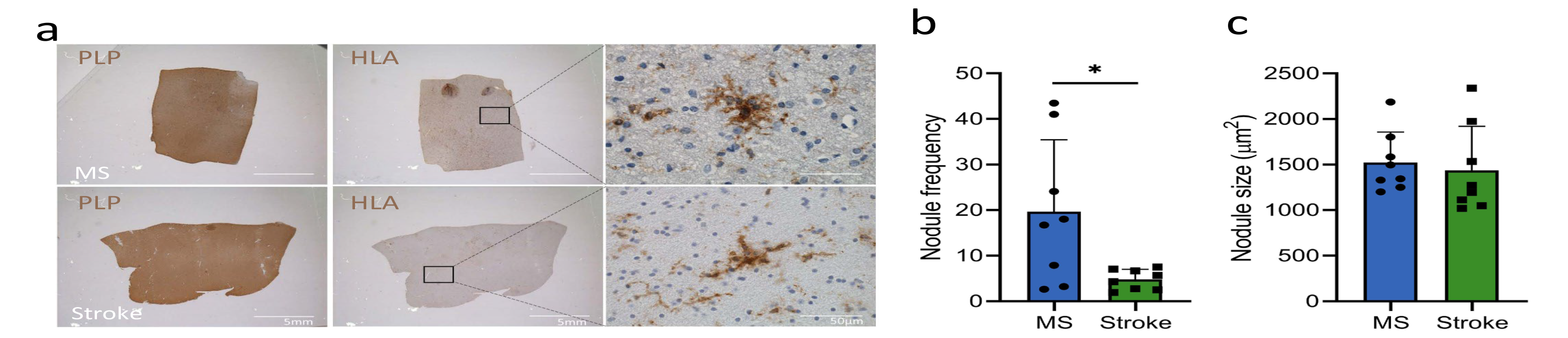
Microglia nodules in MS are more frequent than in stroke, but are similar in size. (**a**) PLP and HLA staining of (NA)WM matter in MS and in stroke shows no sign of demyelination and clustering of HLA-DR^+^ cells into nodules. (**b**) Microglia nodule frequency was calculated as number of microglia nodules per 100 mm^2^. The nodule frequency was higher in MS compared to stroke (p=0.03). (**c**) Microglia nodule size as measured in µm^2^ was similar in MS and stroke. Bar plots show mean ± standard deviation. Significance was tested with a two-sided students t-test, p value <0.05 is indicated with *.

### Gene expression analysis shows diversity between microglia nodules in MS and stroke

To compare gene expression of microglia nodules in MS with those in stroke and non-nodular (NA)WM, tissue was manually dissected using laser capture microscopy. PCA showed partial clustering of four groups distinguishing MS and stroke tissue in the first dimension and nodular and non-nodular (NA)WM tissue in the second dimension (**Fig. 2a**). Cell type deconvolution analysis of the gene expression data showed that microglia nodules in MS compared to MS non-nodular NAWM were characterized by a significantly higher proportion of microglia cells and endothelial cells and a lower proportion of oligodendrocytes. There were no significant differences in estimated cell type proportions between microglia nodules in MS and microglia nodules in stroke, microglia nodules in stroke and stroke non-nodular WM, or MS non-nodular NAWM and stroke non-nodular WM (**Fig. 2b**). As our main comparison of interest was between microglia nodules in MS and stroke, we focused on the results of the analysis without correction for microglia proportion. Results of the analysis with correction for the proportion of microglia are shown in the text (indicated with ‘after correction’) and are also included in **Supplementary Tables 3 and 4**. For each comparison, up to 50 top DE genes are shown in **Supplementary Table 3a-d**. Expression of all genes of interest mentioned below is summarized in **Supplementary Table 4**. The highest number of DE genes were observed in microglia nodules in MS vs MS non-nodular NAWM, where 325 DE genes were upregulated. In microglia nodules in MS vs microglia nodules in stroke, 256 DE genes were upregulated, of which 40 DE genes were also upregulated in microglia nodules in MS vs MS non-nodular NAWM and are considered MS nodule-specific genes. The lowest number of DE genes was found in microglia nodules in stroke vs stroke non-nodular WM with 10 upregulated DE genes. In MS non-nodular NAWM vs stroke non-nodular WM, 23 DE genes were upregulated (**Fig. 2c**). The number of downregulated DE genes was low in all comparisons. Microglia nodules compared to non-nodular (NA)WM in MS and in stroke shared only few communally upregulated DE genes (*C1qB*, *RPGR*, and *SLC11A1),* which are associated with involvement in activation of the classical complement pathway and in phagocytosis. After correction for the microglia proportion (hereafter ‘correction’), *C1qB* remained significantly differentially expressed for both comparisons, while *SLC11A1* was only significantly upregulated by microglia nodules in stroke compared to stroke non-nodular WM (**Fig. 2c-e**).

**Figure 2:**
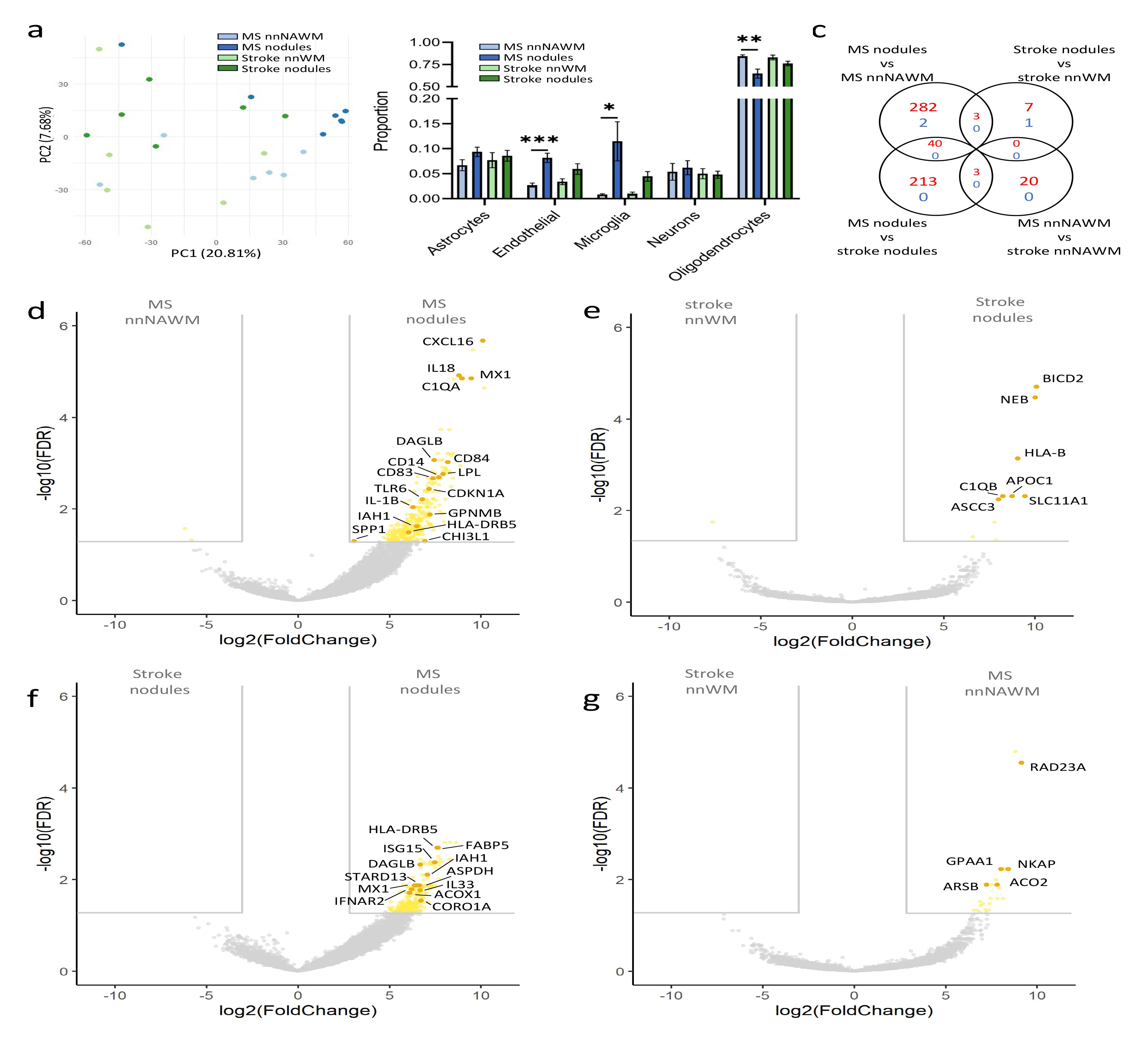
Microglia nodules in MS and stroke share only few commonalities, and microglia nodules in MS show lesion-associated microglia activation. (**a**) PCA plot showing discrimination of MS and stroke samples on the first dimension and discrimination of nodules and non-nodular NAWM in the second component. (**b**) Proportion of cell types in the groups, shown as mean ± standard deviation. Microglia nodules in MS had a higher proportion of endothelial cells and microglia cells, and a lower proportion of oligodendrocytes. p value <0.05 is indicated with *, <0.01 is indicated with **, <0.001 is indicated with ***. (**c**) Venn diagram showing the number of DE genes with adjusted p-value <0.05 and logFC > 3 or <-3, either upregulated in red or downregulated in blue between the various groups. Gene expression of (**d**) MS nodules versus non-nodular NAWM, (**e**) stroke nodules versus stroke non-nodular WM, (**f**) MS nodules versus stroke nodules, and (**g**) MS non-nodular NAWM versus stroke non-nodular WM. DE genes are highlighted in yellow, and top DE genes are highlighted in orange.

### Microglia nodules in MS but not in stroke express MS lesion pathology-associated genes

Microglia nodules compared to non-nodular NAWM in MS had a significantly higher expression of a multitude of genes previously associated with MS pathology (*CXCL16*, *IL18*, *MX1*, *LPL*, *CD14*, *CD83*, *IL1B*, *CDKN1A*, *GPNMB*, *HLA-DRB5*, *C1QA*, *C1qB*, *SPP1*, *TLR6*, *CHI3L1*, after correction *CXCL16*, *IL18*, *C1QA*, *C1qB* remained significant) ^1, 8, 25, 26, 44–50^. *C1qB* was also upregulated by microglia nodules in stroke compared to stroke non-nodular WM, also after correction. Microglia nodules in MS, compared to microglia nodules in stroke, also upregulated expression of genes previously associated with MS-lesion pathology (*HLA-DRB5*, *ISG15*, *MX1*, after correction *HLA-DRB5*, *ISG15*) and MS susceptibility (*IFNAR2*, also after correction) ^45, 51^. In MS non-nodular NAWM compared to stroke non-nodular WM, no genes previously associated with MS pathology were differentially expressed (**Fig. 2d-g**). Thus, microglia nodules in MS and not in stroke show MS lesion-related activation.

### Microglia nodules in MS express genes indicative for lesion formation

Using the DAVID algorithm for functional annotations, we found DE genes in microglia nodules in MS compared to microglia nodules in stroke and compared to MS non-nodular NAWM that functionally may be indicative for lesion formation (**Sup. Fig 1** and **Supplementary Tables 5a-b**). Microglia nodules in MS had upregulated genes that imply involvement in the adaptive and the innate immune response (compared to MS non-nodular NAWM: *HLA-DMB*, *JAK3*, *TLR6*, *IFI27*, none after correction; compared to microglia nodules in stroke: *IDH1*, *PSME3*, *IFNAR2*, *ISG15*, *PARP9*, *IL33*, after correction *IFNAR2*, *ISG15*, *PARP9*), phagocytosis (compared to MS non-nodular NAWM: *CD14*, *IRF8*, *MERTK*, *NCF2*, none after correction), and lipid metabolic processes (compared to MS non-nodular NAWM: *LPL, DAGLB*, *IAH1*, after correction *DAGLB*; compared to microglia nodules in stroke: *PLCD3*, *FABP5*, *ACLY*, *IAH1*, *DAGLB*, *CHI3L1*, *CHI3L2*, *STARD13*, *GPCPD1*, after correction *FABP5*, *IAH1*, *DAGLB*, *STARD13*). Microglia nodules in MS had an increased expression of genes implying T-and B-cell homeostasis and proliferation (compared to MS non-nodular NAWM: *NCKAP1L*, *CASP3*, *JAK3*, *FADD*, *TCIRG1*, after correction *NCKAP1L*; compared to microglia nodules in stroke: *CORO1A*, *NCKAP1L*, *CASP3*, *APBB1IP*, none after correction) and natural killer T (NKT) cell-mediated cytotoxicity (compared to microglia nodules in stroke: *GRB2*, *CASP3*, *BID*, none after correction). In microglia nodules in MS, genes indicating immunoglobulin signaling were upregulated as compared to MS non-nodular NAWM: *HLA-DMA*, *HLA-DPB1*, *HLA-DMB*, *HLA-DRB1*, *HLA-DRB5*, none after correction; compared to microglia nodules in stroke: *HLA-DRB5* (also after correction) and of cytokine signaling, specifically IFNy and TNF (compared to MS non-nodular NAWM: *IRF8*, *CD84*, *LRRK2*, *FADD*, *LPL*, *TLR2*, *EGR1*, *IL18*, after correction *IL18*; compared to microglia nodules in stroke: *IFNAR2,* also after correction). Furthermore, gene expression of microglia nodules in MS indicates they may be under metabolic stress (compared to MS non-nodular NAWM: *PPRC1*, *SAMM50*, *MTG1*, after correction *MTG1*; compared to microglia nodules in stroke: *ACOX1*, *COA4*, *MTG1*, *PPRC1*, *SAMM50*, after correction *MTG1*) and may be responding to as well as producing reactive oxygen species (ROS) (compared to microglia nodules in stroke: *ASPDH*, *BID*, *CRYZL1*, *IDH1*, *MAOB, SMAD3*, none after correction).

### Microglia nodules in MS reside in an inflammatory environment

As gene expression analysis revealed the likelihood of nearby lymphocytes, this was assessed using IHC. In MS and not in stroke, CD20^+^ B cells and CD138^+^ plasma cells were found in close proximity to microglia nodules. CD3^+^ T cells and CD38^+^ plasma blasts were observed more frequently near MS compared to stroke microglia nodules (CD3: MS: 38 of 151 nodules, per donor 26.7% ± 9.1%, stroke: 5 of 89 nodules, per donor 5.1% ± 5.1%, p = 0.02, CD38: MS: 90 of 202 nodules, per donor 34.6% ± 24.5%, stroke: 1 of 65 nodules, per donor 0.4% ± 0.9%, p= 6.7e-3, CD138: MS: 13 of 162 nodules, per donor 6.7% ± 10.7%, stroke: 0 of 78 nodules, per donor 0% ± 0%, **Fig. 3a-e**). In MS, both CD4^+^ as well as CD8^+^ T cells were found in close proximity to microglia nodules in MS (CD4: per donor 8.3 ± 6.8 nodules, CD8: per donor 18.0 ± 16.5 nodules). In MS, subsets of microglia nodules and CD3^+^ T cells expressed the proliferation marker PCNA, suggesting that these T cells have encountered antigenic re-stimulation (**Fig. 3f-j**). In line with the presence of B and plasma cells in close proximity to MS but not near microglia nodules in stroke, we found significantly upregulated expression of the immunoglobulin genes *IGKC*, *IGHG1*, *IGHG2*, and *IGKV3-15* in MS nodule NAWM tissue as compared to stroke (**Fig. 3k**), indicating that immunoglobulin secretion only takes place in tissue containing microglia nodules in MS but not in stroke.

**Figure 3:**
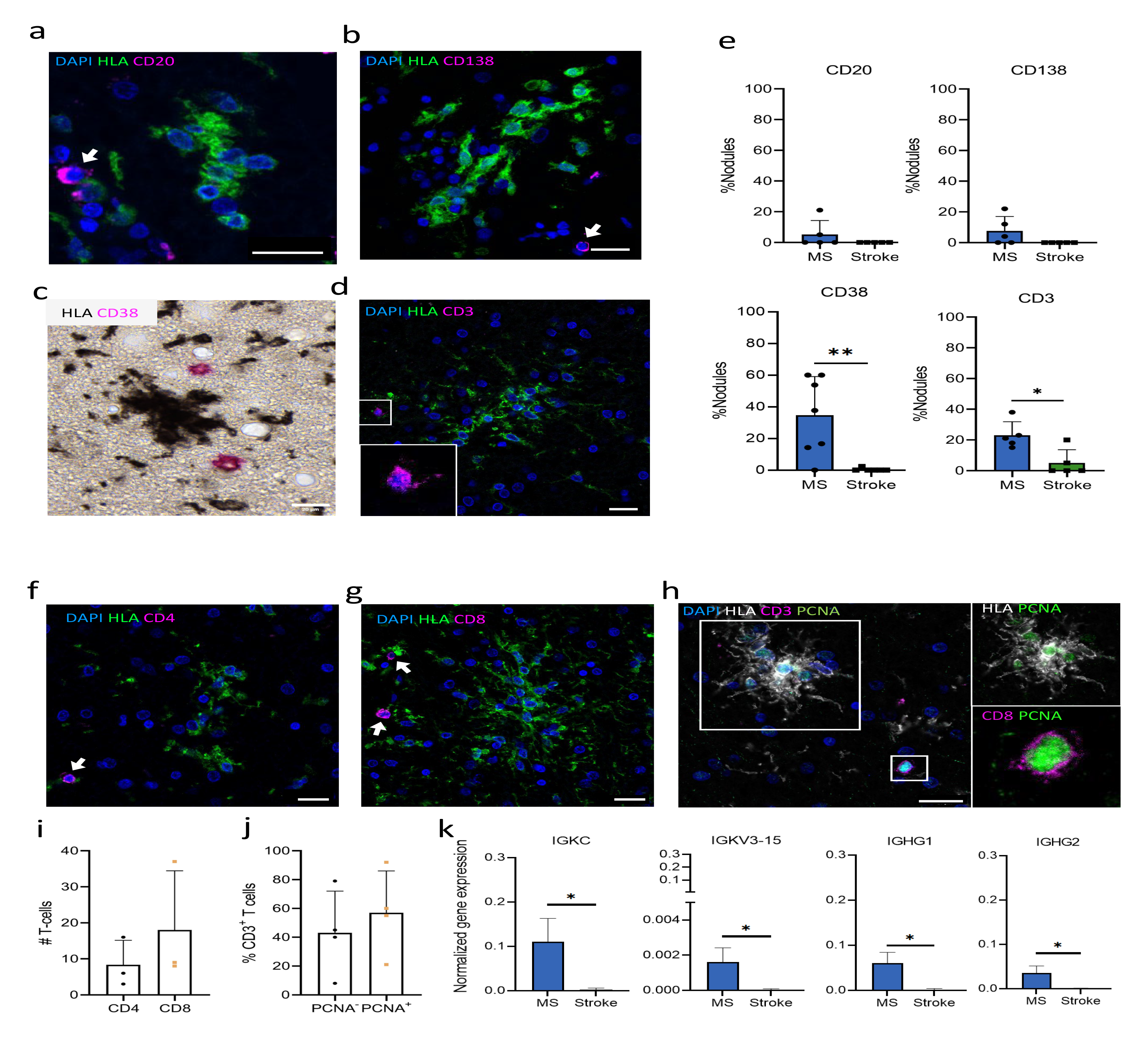
Microglia nodules in MS reside in a more inflammatory environment compared to stroke nodules. Immunohistochemistry of MS tissue showing presence of (a) a CD20^+^ B cell, (b) a CD138^+^ plasma cell, (c) two CD38^+^ plasmablasts, (d) and a CD3^+^ T cell. (e) Quantification of the percentage of nodules in MS and stroke with nearby a CD20^+^ B cell, a CD138^+^ plasma cell, a CD38^+^ plasmablast and a CD3^+^ T cell reveals more B and T cells near microglia nodules in MS than in stroke. Immunohistochemistry showing (f) CD4^+^, (g) CD8^+^, and (h) PCNA^+^ CD3^+^ T cells in close proximity to microglia nodules in MS. Quantification of (i) the number of CD4^+^ and CD8^+^ T cells near microglia nodules in MS and (j) the percentage of PCNA^-^ and PCNA^+^ CD3^+^ T cells near microglia nodules in MS. (k) Immunoglobulin gene expression quantified with RT-qPCR of MS and stroke (NA)WM tissue containing nodules showing higher expression in MS compared to stroke. Bar plots show mean percentage of nodules ± standard deviation. Significance for proportional data was tested with a quasibinomial generalized linear model or a student’s t-test for continuous numerical data, p value <0.05 is indicated with *. Scalebars indicate 20µm.

### Classical complement pathway activation in MS leads to MAC formation

Microglia nodules in MS as well as in stroke have a higher expression of *C1Q* genes compared to non-nodular (NA)WM, which is in line with a previous study showing complement deposition presence in both MS and microglia nodules in stroke ^22^. C1q is a complement component expressed by microglia and macrophages that can bind to the Fc tail of immunoglobulins ^52^ and is a critical mediator of microglia activation in MS ^53^. As immunoglobulin-related genes were only expressed in MS tissue and not in stroke tissue, this may potentially lead to complete activation of the complement cascade causing cell lysis in MS but not in stroke. Therefore, we stained the microglia nodules in MS and in stroke for complement components C1qB, C3d and the membrane attack complex (MAC). In MS and stroke, equal percentages of microglia nodules expressed C1qB (MS: 63 of 163 nodules, per donor 36.6% ± 6.0%, stroke: 22 of 68 nodules, per donor 30.5% ± 19.0%, p=0.25, **Fig. 4a-c**). In MS, microglia nodules more often were associated with C3d^+^ axons compared to microglia nodules in stroke (MS: 35 of 150 nodules, per donor 17.2% ± 15.5%, stroke: 4 of 54 nodules, per donor 4.2% ± 7.0%, p=0.008), suggesting an increased activation of the complement pathway in MS (**Fig. 4d-f**). Furthermore, microglia nodules in MS were more often associated with C5b-9, which constitutes the MAC, compared to microglia nodules in stroke (MS: 29 of 106 nodules, per donor 31.40% ± 17.37, stroke: 16 of 117 nodules, per donor 12.96% ± 7.54%, p=0.03), indicating involvement of microglia nodules in MS in complement mediated tissue degeneration (**Fig. 4g-i**).

**Figure 4:**
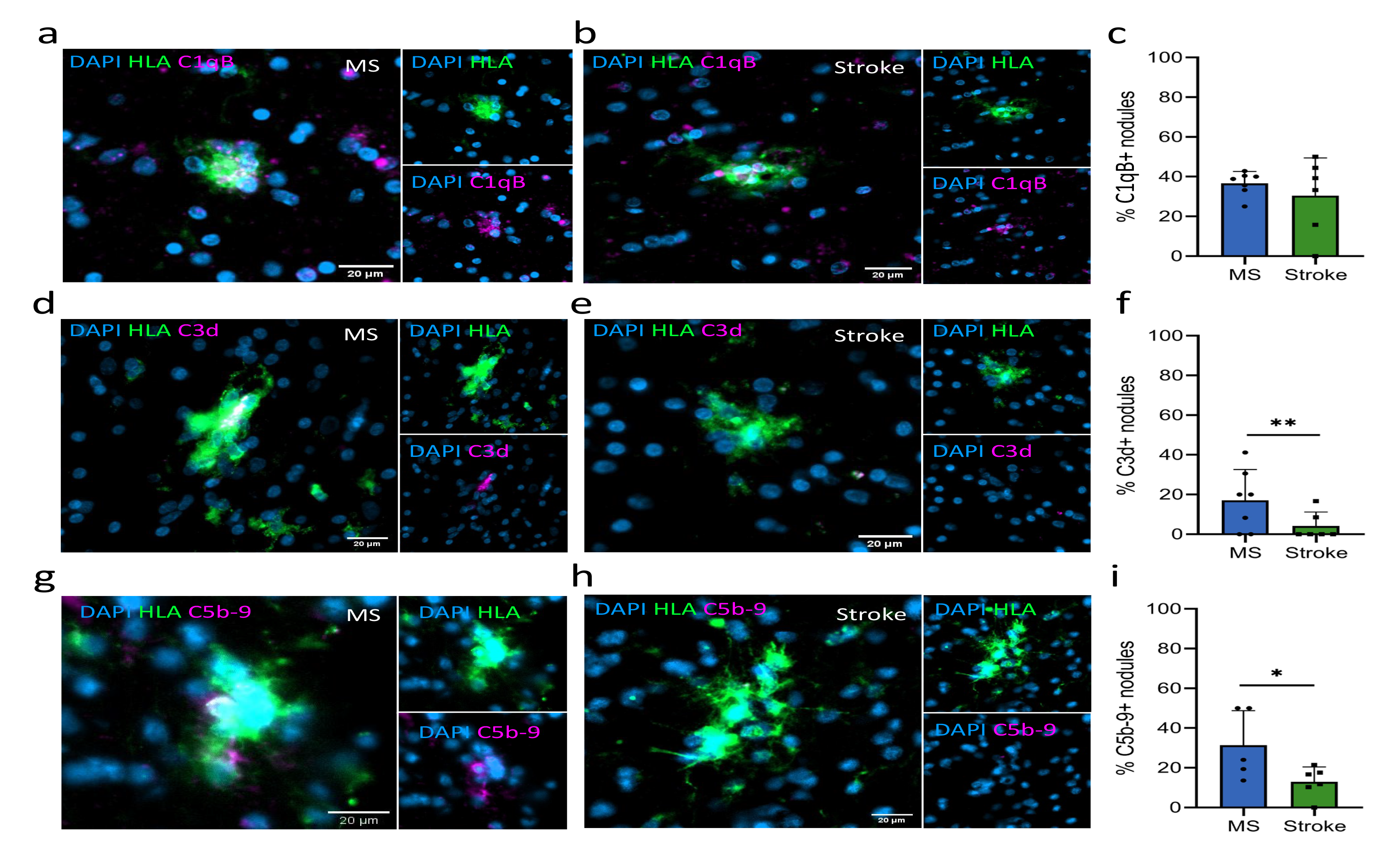
Microglia nodules in MS are associated with activation of the classical complement pathway and are associated with membrane attack complex formation. Immunohistochemistry of a C1qB^+^ HLA^+^ microglia nodule (a) in MS and (b) in stroke. (c) Microglia nodules in MS and in stroke equally often express C1qB. Immunohistochemistry of (d) a HLA^+^ microglia nodule associated with a C3d^+^ axonal fragment in MS and (e) an HLA^+^ microglia nodule in stroke that is not associated with a C3d^+^ axonal fragment. (f) In MS, microglia nodules are more often associated with C3d^+^ axonal fragments than in stroke. Immunohistochemistry of (g) an HLA^+^ C5b-9^+^ microglia nodule in MS and (h) an HLA^+^ microglia nodule in stroke that is not C5b-9^+^. (i) In MS, microglia nodules are more often C5b- 9^+^ than in stroke. Bar plots show mean value ± standard deviation. Significance was tested with a quasibinomial generalized linear model, p value <0.05 is indicated with *, p value <0.01 is indicated with **.

### Increased lipid metabolism in microglia nodules in MS

Genes of interest involved in lipid metabolism that were upregulated in microglia nodules in MS compared to microglia nodules in stroke were validated with IHC. Microglia nodules in MS compared to microglia nodules in stroke significantly more often expressed fatty acid-binding protein 5 (FABP5) (MS: 81 of 143 nodules, per donor 55.2% ± 8.7%, stroke: 14 of 56 nodules, per donor 22.0% ± 19.6%, p=3.3^-6^), diacylglycerol lipase-beta (DAGLB) (MS: 62 of 79 nodules, per donor 73.8% ± 10.0%, stroke: 12 of 35 nodules, per donor 19.2% ± 22.5%, p=1.11^-8^), and StAR-related lipid transfer domain protein 13 (STARD13) (MS: 28 of 57 nodules, per donor 47.5% ± 19.9%, stroke: 7 of 24 nodules, per donor 25.1% ± 20.9%, p<0.05). Difference in expression of isoamyl acetate hydrolyzing esterase 1 (IAH1) did not reach significance due to large variation (MS: 21 of 88 nodules, per donor 43.7% ± 31.0%, stroke: 7 of 24 nodules, per donor 25.1% ± 20.9%, p=0.32, **Fig. 5a-l**).

**Figure 5:**
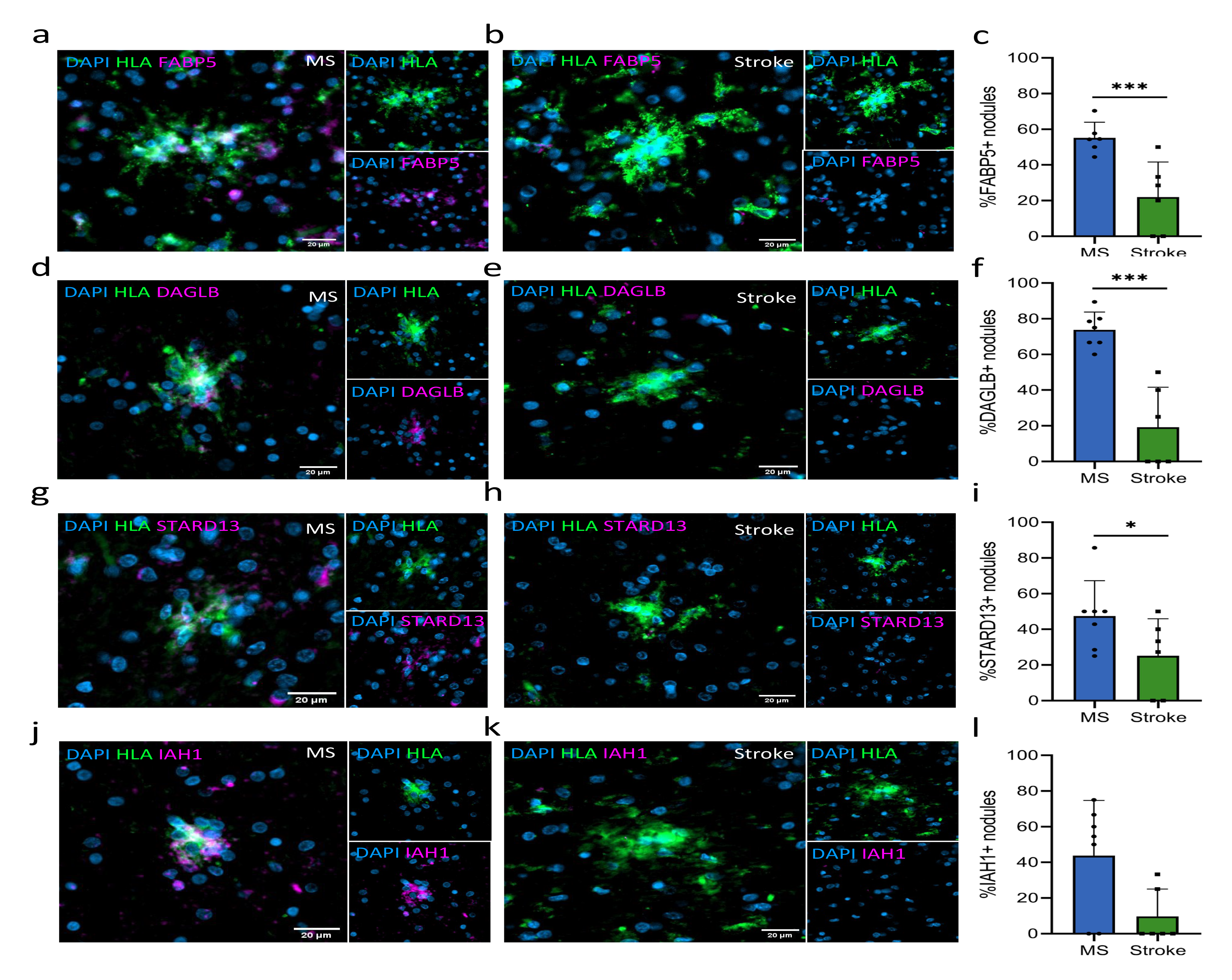
Microglia nodules in MS are involved in lipid metabolism. Immunohistochemistry of (a) an HLA^+^ FABP5^+^ microglia nodule in MS and (b) an HLA^+^ FABP5^-^ microglia nodule in stroke. (c) In MS, microglia nodules more often express FABP5 than in stroke. Immunohistochemistry of (d) an HLA^+^ DAGLB^+^ microglia nodule in MS and (e) an HLA^+^ DAGLB^-^ microglia nodule in stroke. (f) In MS, microglia nodules more often express DAGLB compared to microglia nodules in stroke. Immunohistochemistry of (g) an HLA^+^ STARD13^+^ microglia nodule in MS and (h) an HLA^+^ STARD13^-^ microglia nodule in stroke. (i) In MS, microglia nodules more often express STARD13 compared to microglia nodules in stroke. Immunohistochemistry of (j) an HLA^+^ IAH1^+^ microglia nodule in MS and (k) an HLA^+^ IAH1^-^ microglia nodule in stroke. (l) Microglia nodules in MS do not significantly more often express IAH1 compared to those in stroke. Bar plots show mean value ± standard deviation. Significance was tested with a quasibinomial generalized linear model, p value <0.05 is indicated with *, <0.01 is indicated with **, <0.001 is indicated with ***. Bar plots show mean value ± standard deviation.

In MS, the NAWM has more oxidized phospholipids compared to controls ^54^, which may be one of the triggers for microglia in MS to cluster and form nodules in order to clear up the oxidized phospholipids. Therefore, using IHC and STED microscopy, the percentage of microglia nodules that had phagocytosed oxidized phospholipids were quantified in MS and stroke. In microglia nodules in MS, lysosomal-associated membrane protein 1 (Lamp1)^+^ lysosomes more often contained oxidized phospholipids (detected using antibody E06) compared to microglia nodules in stroke (MS: 30 of 47 nodules, per donor 68.0% ± 2.8%, stroke: 7 of 29 nodules, per donor 29.1% ± 7.0%, p=3.2^-12^), showing that microglia nodules in MS are more involved in phagocytosis of oxidized phospholipids (**Fig. 6a-c**).

**Figure 6:**
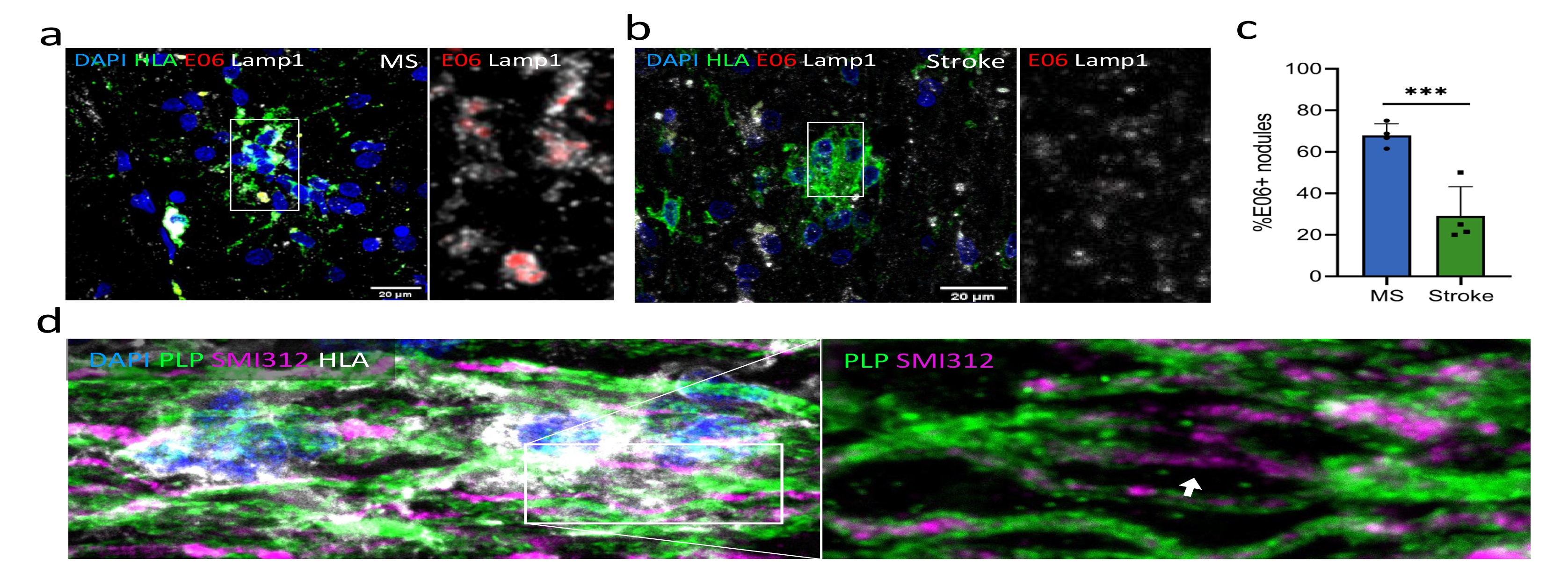
Microglia nodules in MS are involved in demyelination. STED microscopy of (a) a microglia nodule in MS containing lysosomal (Lamp1) oxidized phospholipids (E06) and (b) a microglia nodule in stroke that does not show oxidized phospholipids (E06) in the lysosomes (Lamp1). (c) Microglia nodules in MS have more often phagocytosed oxidized phospholipids compared to microglia nodules in stroke. Bar plot shows mean value ± standard deviation. Significance was tested with a quasibinomial generalized linear model, p value <0.001 is indicated with ***. (d) In MS NAWM in the optic nerve, partially demyelinated axons (PLP^-^) encapsulated by microglial nodules were found.

As microglia nodules in MS are likely involved in lipid metabolism, we set out to investigate if microglia nodules are involved in demyelination with high-resolution IHC. Cryo-protected MS normal-appearing optic nerve tissue ^29^ provided sufficient resolution to investigate demyelination of individual axons encapsulated by microglia nodules. Some axons (42%) surrounded by microglia nodules showed partial demyelination, as indicated by loss of PLP staining of a part of the SMI312^+^ axon (**Fig. 6d**). Therefore, we conclude that microglia nodules in MS are indeed involved in demyelination, and a subset of nodules are forming a ‘mini lesion’.

### Mitochondrial network in microglia nodules in MS is more tubular

DE genes of microglia nodules in MS compared to microglia nodules in stroke were indicative of an altered metabolic state. Using IHC and STED microscopy, we assessed the mitochondrial network of microglia nodules in MS and in stroke. Of each nodule, the translocase of the outer mitochondrial membrane (TOMM20) network was classified as fragmented, intermediate, or tubular. Quantification of the mitochondrial network in microglia nodules in MS and in stroke showed that in MS the mitochondrial network was more often tubular (MS: 51.0% ± 18.5%, Stroke: 13.1% ± 14.9%, p=0.009) (**Fig. 7a-c**). This tubular network is indicative of a hypermetabolic state of the microglia nodules in MS.

**Figure 7:**
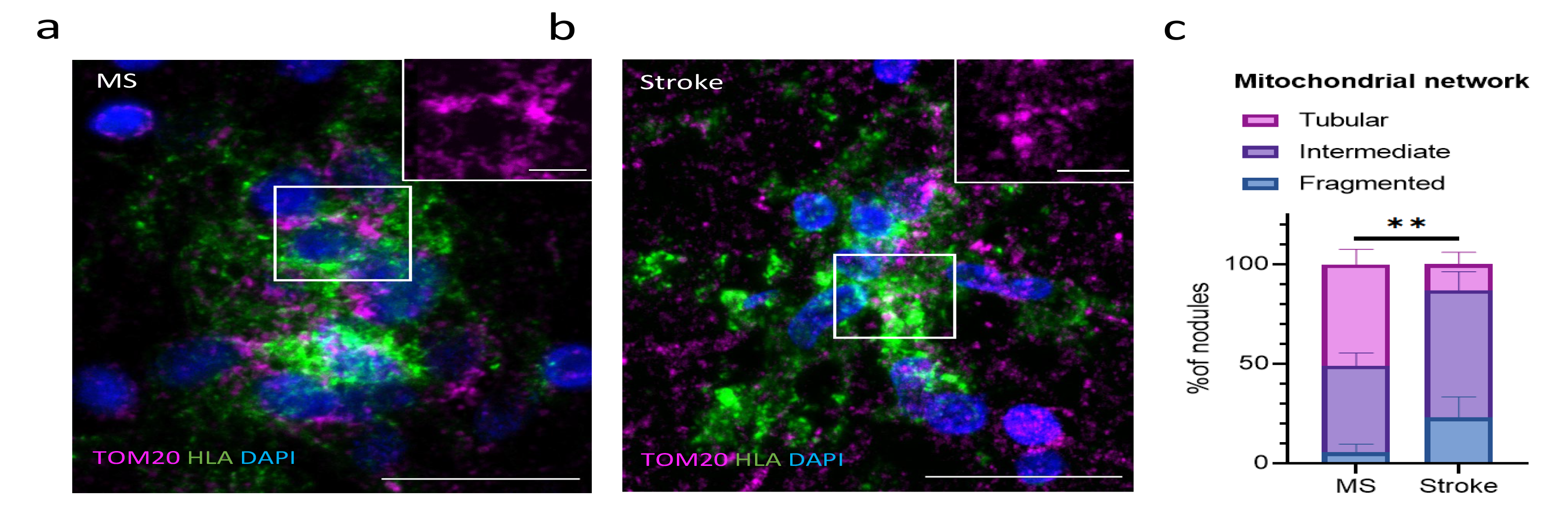
Microglia nodules in MS show a more tubular mitochondrial network. STED microscopy of (a) an HLA^+^ microglia nodule in MS with a fu ed and tubular TOM20 mitochondrial network and (b) an HLA^+^ microglia nodule in stroke with a fragmented TOM20 mitochondrial network. (c) Microglia nodules in MS more often showed a tubular and fused mitochondrial network compared to microglia nodules in stroke. Bar plot showing mean percentage per type of mitochondrial network in MS and in stroke. Significance was tested with a quasibinomial linear model, p value <0.01 is indicated with **.

### Data availability

The RNA-sequencing dataset will be made available after publication in the Gene Expression Omnibus (GEO) database.

## Discussion

Here, we have studied the potential involvement of microglia nodules in MS-lesion formation by correlating their presence with pathological and clinical characteristics and by comparing microglia nodules in MS to microglia nodules in stroke and to surrounding non-nodular (NA)WM using IHC and RNA sequencing of laser-microdissection captured tissue. We show that MS microglia nodules (1) correlate with more severe MS pathology, (2) upregulate expression of genes similar to MS-lesions, (3) upregulate expression of genes associated with adaptive and innate immune responses, lymphocyte activation, phagocytosis, lipid metabolism, and metabolic stress, (4) have nearby lymphocyte presence, (5) contain partially demyelinated axons, (6) are associated with IgG-transcription, complement activation, and MAC formation, and (7) are in a hypermetabolic state associated with increased pro-inflammatory cytokine and ROS gene expression. Our findings indicate that the nodule milieu may be held responsible for early lesion formation.

We here show that microglia nodules in MS are pathologically relevant. MS donors with microglia nodules compared to those without microglia nodules had a higher lesion load and reactive site load. The proportion of active lesions was higher, and the proportion of inactive and remyelinated lesions was lower in MS donors with microglia nodules compared to those without. Interestingly, there was no difference in the proportion of mixed active/inactive lesions. As we hypothesize that active lesions precede mixed active/inactive lesions, this indicates that microglia nodules are possibly associated with new MS lesion formation. Therefore, MS donors without microglia nodules may represent a subgroup in which frequency of new lesion formation has decreased, whereas those with microglia nodules may still develop new MS lesions. Presence of microglia nodules was not associated with clinically more severe MS. Although microglia nodules in MS do not seem to represent previous clinical progression, this does not exclude the possibility that they may represent future clinical progression. Furthermore, clinical progression is associated with more factors than new lesion formation, such as atrophy and ongoing axonal damage ^55^.

Nodule tissue in MS compared to NAWM contained a higher proportion of microglia, a lower proportion of oligodendrocytes, and a higher proportion of endothelial cells. The lower proportion of oligodendrocytes may indicate demyelination of axons encapsulated by nodules. Vessels are preferential lesion formation sites ^56^, and previously we have shown that in the NAWM, there is an increase in perivascular tissue-resident memory T cells, which may be influencing the milieu in which microglia nodules in MS reside ^57^. Therefore, the cell type composition of nodule tissue in MS suggests that microglia nodules may represent the first stage of MS lesion formation.

Microglia nodules highly expressed genes that were previously implicated in MS pathology, such as *LPL*, *CXCL16*, *CD14*, *CDKN1A*, and *CHI3L1* ^8, 48^. Moreover, *CXCL16*, *MX1*, *HLA-DRB5*, *ISG15*, *IFNAR2*, and *IL1B* were previously found upregulated in active lesions or the rim of mixed active/inactive lesions, and *CXCL16* and *CHI3L1*, related to lipid binding, were found upregulated in perilesional areas of active lesions and are indicative for the expansion of lesions ^25, 26, 45^. *CXCL16*, *CHI3L1*, *IFNAR2*, and *HLA-DRB5* have furthermore been suggested as potential prognostic markers in MS ^46, 49–51^. From this, we conclude that microglia nodules in MS show signs of lesion-associated microglia activation and are not part of a diffuse reaction of chronic damage in MS NAWM.

Interestingly, we found MS microglia nodule-specific upregulation of genes associated with lipid metabolism and catabolism (*DAGLB, IAH1, PLCD3, FABP5, ACLY, CHI3L2, STARD13*, and *GPCPD1*)^43^. On the protein level, microglia nodules in MS compared to in stroke were more often FABP5^+^, DAGLB^+^, or STARD13^+^. However, the percentage of IAH1^+^ microglia nodules was only numerically higher in MS compared to stroke, likely indicating that the increased expression of IAH1 is also driven by other cell types than by microglia alone. We hypothesize that lipid metabolic processes are key in progression of an MS nodule to an inflammatory demyelinating lesion. Previously, we have shown that there are more mitochondria in axons in the NAWM ^29^ that may precede lipid oxidation in MS ^54^. Potentially, this is an important trigger in the formation of microglia nodules. Therefore, microglia nodules in MS and in stroke were stained for lysosomal oxidized phospholipids. Indeed, in MS, a higher percentage of microglia nodules contained phagocytosed oxidized lipids, indicating that potentially these microglia nodules had formed to clear up damaged myelin. Interestingly, in normal-appearing optic nerve tissue of MS donors, some axons encapsulated by microglia nodules were partially demyelinated. Considering the differential gene expression and the partially demyelinated axons in the nodules in MS, some nodules seem to be involved in demyelination and the formation of new lesions.

In contrast to what was previously found ^22^, with gene expression analysis and IHC we show the presence of C1q associated with microglia nodules in both MS and stroke. As complement deposition is necessary for Wallerian degeneration ^58^, and Wallerian degeneration is a commonality between MS and stroke, microglia nodules in MS and stroke were likely to possess similarities.

Gene expression analysis indicates close proximity of activated lymphocytes that are influencing and being influenced by the microglia nodules in MS and not in stroke (*NCKAP1L, CASP3, JAK3, TCIRG1, CORO1A, GRB2, IRF8, TLR2, IL18)*. With IHC, we found CD20^+^ B cells and CD138^+^ plasma cells in close proximity to microglia nodules in MS and not in stroke, and in MS more microglia nodules were associated with activated, proliferating CD3^+^ T cells and CD38^+^ plasma blasts. Previously, Van Noort *et al*. (2011) have hypothesized that microglia nodules in MS no longer reflect a local neuroprotective and reparative response if there is presence of lymphocytes. Possibly, the presence of activated T cells and immunoglobulin-producing B-cell blasts near some microglia nodules together with the phagocytosis of oxidized lipids in MS creates a volatile situation. This might indicate a critical turning point in which the nodule is not able to resolve and progresses into a demyelinating and inflammatory site. Previously, we showed MS NAWM to be enriched for perivascular B cells and T cells ^57, 59^. These lymphocytes may produce soluble factors contributing to lesion formation as cytokines and immunoglobulins ^44, 60, 61^. Therefore, as we show lymphocytes in close proximity to microglia nodules in MS, they may be contributing to the inflammatory environment in which microglia nodules reside.

Moreover, we found that tissue containing microglia nodules in MS had significant upregulated immunoglobulin genes (*IGKC*, *IGHG1*, *IGHG2*, and *IGKV3-15*), which was absent in tissue containing stroke nodules. This corresponds with the characteristic high prevalence of intrathecal unique oligoclonal IgG production in MS and the presence of B cells in MS NAWM ^61^. Previously, we have shown that microglia in the NAWM are immunosuppressed ^62^, and that immunoglobulins can break this immune tolerance of microglia cells through Fcγ receptors and thereby potentiate inflammation by microglia ^63^. Therefore, the presence of immunoglobulin-producing cells near microglia nodules in MS in combination with the activation of microglia through pro-inflammatory cytokines secreted by nearby, activated lymphocytes may represent a hazardous situation. Furthermore, complement deposition was found in microglia nodules in both stroke as well as in MS, but only in MS immunoglobulins were found. This may lead to activation of the complement cascade and subsequently to MAC formation and cell death in MS and not in stroke ^64^. Previously, it was shown that microglia nodules, both in stroke as well as in MS, encapsulate C3d^+^ complement deposits associated with the degeneration of axons ^15, 22, 27^. Here, we demonstrate that microglia nodules in MS are more often associated with C3d^+^ complement deposits compared to microglia nodules in stroke, and microglia nodules in MS were also more often associated with MAC formation. This may be a key mechanism underlying progression of microglia nodules in MS to immunoglobulin-associated, complement-dependent, demyelinating inflammatory lesions. These findings suggest that MS therapies associated with loss of intrathecal oligoclonal bands may also associate with a reduced odds of nodules progressing towards a lesion. Likewise, a higher intrathecal oligoclonal immunoglobulin-production at the first symptoms of MS is associated with higher odds of new lesion development on MRI and a swifter occurrence of relapses ^65–67^.

Our data indicate that microglia nodules in MS have a disturbed cell metabolism (*PPRC1*, *SAMM50*, *MTG1*, *ACOX1*, *COA4*) and are responding to as well as producing ROS (*ASPDH*, *BID*, *CRYZL1*, *IDH1*, *MAOB*, *SMAD3*). Correction for the proportion of microglia in the MS nodules negated significance, however this is not very surprising as microglia are main producers of ROS and therefore, the proportion of microglia cells may play a role in the neurodegenerative properties of an MS nodule. Generally, activated microglia cells switch from oxidative phosphorylation to glycolysis and form a fragmented mitochondria network ^68^. Notably, microglia nodules in MS generally possess a more tubular and less fragmented mitochondrial network compared to microglia nodules in stroke. We hypothesize that the combined activation of microglia cells by surrounding lymphocytes together with the phagocytosis of oxidized lipids may result in a hypermetabolic and hyperinflammatory state, as previously shown for atherosclerosis ^69, 70^, in which the microglia rely both on glycolysis as well as oxidative phosphorylation. This can result in prolonged longevity and increased production of cytokines and ROS. We further show that microglia nodules in MS compared to stroke have upregulated genes associated with production of cytokines, specifically IFNy and TNF (*IRF8*, *GADD*, *IFNAR2*, *CD84*, *LRRK2*, *FADD*, *LPL*, *TLR2*, *EGR1*, and *IL18*). Future studies need to elucidate a causal relation between the combined stimulation of the microglia cells through pro-inflammatory cytokines and phagocytosis of oxidized phospholipids, the shift in mitochondrial network, and the production of cytokines, as this opens up a potentially interesting therapeutic avenue.

In summary, we propose that some microglia nodules in MS that have the potential to progress into inflammatory and demyelinating MS lesions, whereas those in stroke will not. Therefore, differences between microglia nodules in MS and stroke can provide insight in mechanisms behind MS lesion formation. Here, we demonstrate that microglia nodules in MS upregulate lesion-associated genes and genes indicative for demyelination. We also show that some microglia nodules in MS encapsulate partially demyelinated axons. Moreover, we here describe that a combination of activated T cells, immunoglobulin-producing B cells, and oxidized lipids in and around MS microglia nodules may together enable microglia nodules to become hypermetabolic, form MACs, and give rise to ‘mini’ MS lesions. Together, we conclude that microglia nodules in MS are likely sites of lesion initiation and represent an interesting therapeutic target to prevent early demyelination and MS lesion formation.

## Author’s contribution

A.M.R.v.d.B., M.v.d.P, J.S, I.H., and J.H. contributed to conception and design of the study, A.M.R.v.d.B, M.v.d.P., and M.C.J.V. contributed to collection of sequencing samples, A.M.R.v.d.B, A.J., H.J.E., and P.D.M. contributed to analysis of sequencing results, A.M.R.v.d.B., M.v.d.P., N.F., and A.M.B. contributed to acquisition and analysis of pathological data. All authors contributed to interpretation of findings and drafting and editing of the manuscript.

## Acknowledgements

We are grateful to the brain donors and their families for their commitment to the Netherlands Brain Bank donor program. Funding for this research was obtained from MS Research grants 17-975 and 19- 1079.

---

**Supplementary figure 1:**
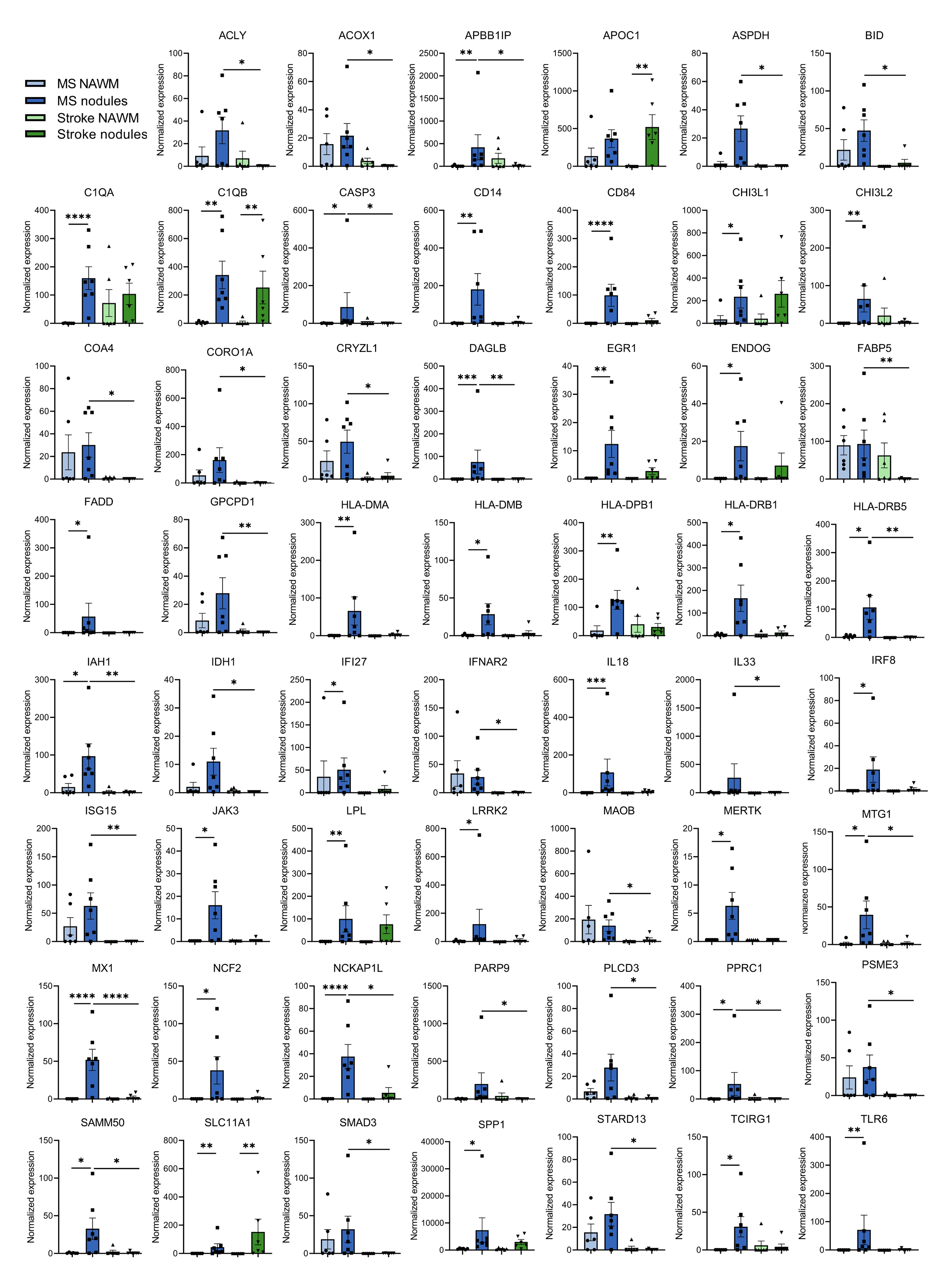
Normalized expression of genes likely involved in functional pathways relevant for MS pathology. Data is shown as mean ± standard deviation. Normalized gene expression is calculated as 2^log(CPM)^. p value <0.05 is indicated with *, <0.01 is indicated with **, <0.001 is indicated with ***.

**Supplementary table 1:** Outliers removed from RNA sequencing analysis.

| Sequence ID |  | Diagnosis | Tissue | Age | Gender |
| --- | --- | --- | --- | --- | --- |
| s104343_001_003 | 94-116 | Stroke | Microglia nodules | 72 | F |
| s104343_001_004 | 94-116 | Stroke | NAWM | 72 | F |
| s104343_001_028 | 17-100 | MS | NAWM | 66 | M |

**Supplementary table 2:**
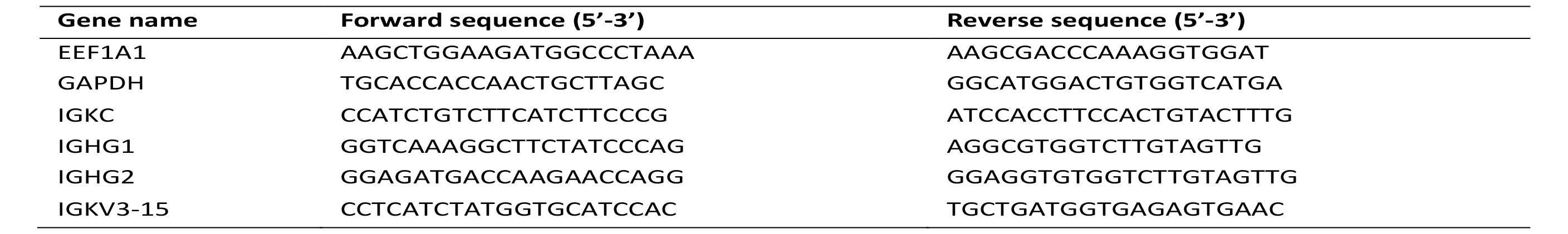
Primers used for RT-qPCR on nodule tissue.

**Supplementary table 3a:** Top 50 upregulated DE genes in MS nodules vs MS nnNAWM.

| Gene symbol | Gene name | logFC | P. Adj. | LogFC cor. | P. Adj. Cor. |
| --- | --- | --- | --- | --- | --- |
| RGS16 | regulator of G protein signaling 16 | 10.186 | 2.27e-5 | 9.149 | 0.015 |
| CXCL16 | C-X-C motif chemokine ligand 16 | 10.074 | 2.11e-6 | 9.666 | 0.004 |
| TNFRSF1B | TNF receptor superfamily member 1B | 9.578 | 3.36e-6 | 9.079 | 0.006 |
| C1QA | complement C1q A chain | 9.451 | 1.40e-5 | 8.344 | 0.022 |
| CTSC | cathepsin C | 9.025 | 1.40e-5 | 7.611 | 0.033 |
| MX1 | MX dynamin like GTPase 1 | 8.935 | 1.40e-5 | 8.283 | 0.022 |
| RGS1 | regulator of G protein signaling 1 | 8.869 | 0.001 | 7.142 | 0.162 |
| EML4-AS1 | EML4 antisense RNA 1 | 8.846 | 1.21e-5 | 8.333 | 0.015 |
| IL18 | interleukin 18 | 8.773 | 1.21e-5 | 8.771 | 0.008 |
| NCKAP1L | NCK associated protein 1 like | 8.491 | 1.46e-5 | 7.309 | 0.033 |
| SRP68 | signal recognition particle 68 | 8.479 | 0.001 | 9.201 | 0.026 |
| SLC7A5 | solute carrier family 7 member 5 | 8.454 | 0.002 | 9.215 | 0.026 |
| LCP1 | lymphocyte cytosolic protein 1 | 8.352 | 0.001 | 7.297 | 0.162 |
| APBB1IP | amyloid beta precursor protein binding family B member 1 interacting protein | 8.350 | 0.003 | 7.614 | 0.147 |
| C1QB | complement C1q B chain | 8.315 | 0.001 | 7.831 | 0.037 |
| ERCC5 | ERCC excision repair 5, endonuclease | 8.267 | 1.85e-4 | 8.427 | 0.026 |
| SLCO2B1 | solute carrier organic anion transporter family member 2B1 | 8.257 | 0.002 | 7.878 | 0.051 |
| IGSF6 | immunoglobulin superfamily member 6 | 8.171 | 0.001 | 7.135 | 0.111 |
| CD84 | CD84 molecule | 8.169 | 0.001 | 5.701 | 0.342 |
| WARS1 | tryptophanyl-tRNA synthetase 1 | 8.121 | 0.002 | 7.574 | 0.156 |
| VAMP8 | vesicle associated membrane protein 8 | 8.111 | 0.001 | 6.343 | 0.270 |
| XAF1 | XIAP associated factor 1 | 8.090 | 0.001 | 7.407 | 0.156 |
| SAT2 | spermidine/spermine N1-acetyltransferase family member 2 | 8.049 | 0.006 | 8.251 | 0.153 |
| SCIN | scinderin | 7.971 | 0.019 | 6.159 | 0.402 |
| GPRIN3 | GPRIN family member 3 | 7.928 | 0.001 | 7.268 | 0.147 |
| MED11 | mediator complex subunit 11 | 7.922 | 0.001 | 7.171 | 0.170 |
| LPL | lipoprotein lipase | 7.916 | 0.002 | 5.355 | 0.395 |
| FCER1G | Fc fragment of IgE receptor Ig | 7.897 | 0.013 | 6.426 | 0.347 |
| CAMKK2 | calcium/calmodulin dependent protein kinase kinase 2 | 7.877 | 0.002 | 6.950 | 0.200 |
| RPGR | retinitis pigmentosa GTPase regulator | 7.790 | 1.85e-4 | 7.929 | 0.029 |
| ARHGAP24 | Rho GTPase activating protein 24 | 7.787 | 0.002 | 6.099 | 0.321 |
| ARHGAP18 | Rho GTPase activating protein 18 | 7.783 | 0.001 | 6.861 | 0.147 |
| MRPL55 | mitochondrial ribosomal protein L55 | 7.750 | 0.011 | 6.202 | 0.453 |
| HLA-DPB1 | major histocompatibility complex, class II, DP beta 1 | 7.735 | 0.008 | 7.106 | 0.203 |
| PLEK | pleckstrin | 7.697 | 0.003 | 5.824 | 0.321 |
| CD14 | CD14 molecule | 7.690 | 0.002 | 6.245 | 0.231 |
| LPAR6 | lysophosphatidic acid receptor 6 | 7.646 | 0.001 | 7.031 | 0.051 |
| EML4 | EMAP like 4 | 7.638 | 0.004 | 7.184 | 0.147 |
| TRIM52 | tripartite motif containing 52 | 7.613 | 0.002 | 8.153 | 0.070 |
| SERPINA1 | serpin family A member 1 | 7.551 | 0.006 | 6.582 | 0.353 |
| SLC2A5 | solute carrier family 2 member 5 | 7.542 | 0.002 | 5.592 | 0.373 |
| MPPE1 | metallophosphoesterase 1 | 7.492 | 0.002 | 7.091 | 0.164 |
| ETV5 | ETS variant transcription factor 5 | 7.488 | 0.003 | 7.152 | 0.188 |
| TANC1 | tetratricopeptide repeat, ankyrin repeat and coiled-coil containing 1 | 7.484 | 0.009 | 7.899 | 0.093 |
| DAGLB | diacylglycerol lipase beta | 7.432 | 0.001 | 7.622 | 0.053 |
| ITGAX | integrin subunit alpha X | 7.412 | 0.006 | 6.118 | 0.352 |
| LAPTM5 | lysosomal protein transmembrane 5 | 7.410 | 0.029 | 6.363 | 0.334 |
| RCOR1 | REST corepressor 1 | 7.404 | 0.006 | 5.679 | 0.407 |
| AXL | AXL receptor tyrosine kinase | 7.396 | 0.003 | 6.047 | 0.342 |

**Supplementary table 3b:** Upregulated DE genes in stroke nodules vs stroke nnNAWM.

| Gene symbol | description | logFC | P. Adj. | LogFC cor. | P. Adj. Cor. |
| --- | --- | --- | --- | --- | --- |
| BICD2 | BICD cargo adaptor 2 | 10.061 | 1.97e-5 | 9.803 | 2.85e-4 |
| NEB | nebulin | 9.979 | 3.37e-5 | 9.785 | 4.50e-4 |
| HLA-B | major histocompatibility complex, class I, B | 9.439 | 0.005 | 9.061 | 0.010 |
| C1QB | complement C1q B chain | 9.042 | 0.001 | 8.891 | 0.002 |
| APOC1 | apolipoprotein C1 | 8.731 | 0.005 | 8.497 | 0.009 |
| SLC11A1 | solute carrier family 11 member 1 | 8.227 | 0.005 | 7.755 | 0.010 |
| ASCC3 | activating signal cointegrator 1 complex subunit 3 | 7.989 | 0.006 | 8.115 | 0.013 |
| DIS3L2 | DIS3 like 3'-5' exoribonuclease 2 | 7.840 | 0.044 | 8.301 | 0.072 |
| WASL | WASP like actin nucleation promoting factor | 7.779 | 0.018 | 7.885 | 0.033 |
| RPGR | retinitis pigmentosa GTPase regulator | 6.599 | 0.037 | 6.644 | 0.073 |

**Supplementary table 3c:**
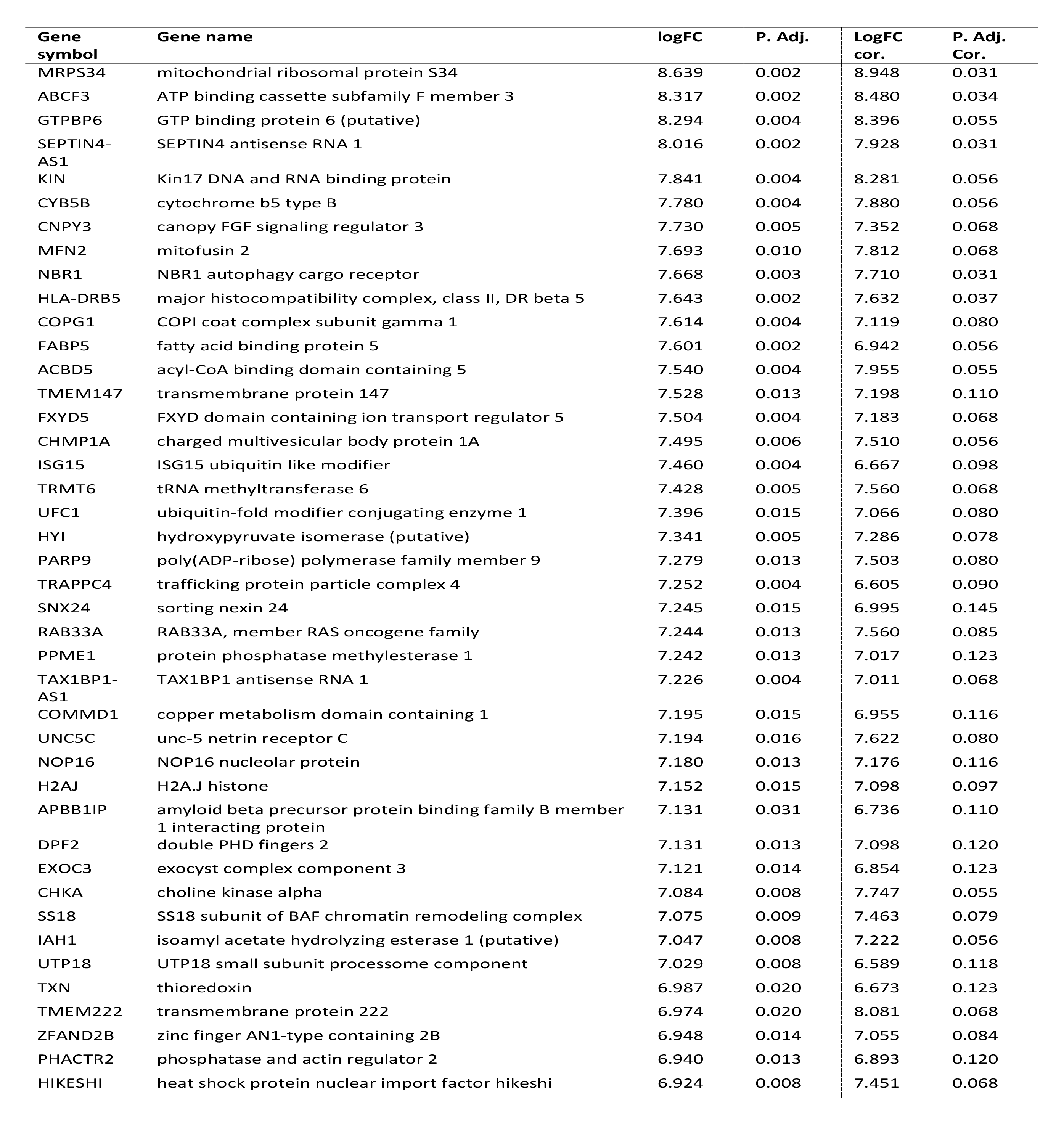
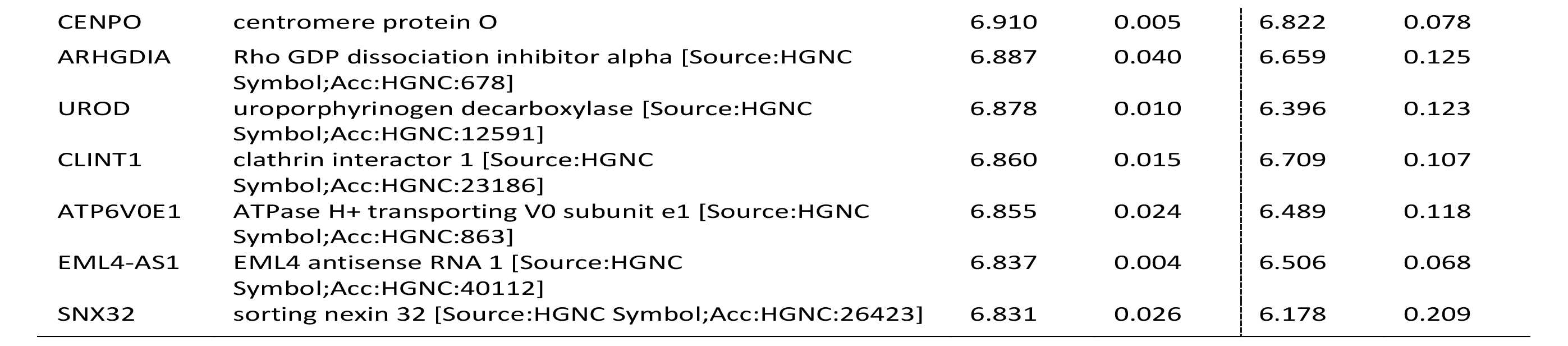
Top 50 upregulated DE genes in MS nodules vs stroke nodules.

**Supplementary table 3d:** Upregulated DE genes in MS nnNAWM vs stroke nnNAWM.

| Gene symbol | Gene name | logFC | P. Adj. | LogFC cor. | P. Adj. Cor. |
| --- | --- | --- | --- | --- | --- |
| RAD23A | RAD23 homolog A, nucleotide excision repair protein | 9.132 | 2.92e-5 | 9.139 | 3.78e-5 |
| NSUN5 | NOP2/Sun RNA methyltransferase 5 | 8.827 | 1.61e-5 | 8.825 | 2.75e-5 |
| COPA | COPI coat complex subunit alpha | 8.435 | 0.006 | 8.435 | 0.008 |
| NKAP | NFKB activating protein | 8.405 | 0.006 | 8.401 | 0.008 |
| ARFGAP2 | ADP ribosylation factor GTPase activating protein 2 | 8.202 | 0.006 | 8.219 | 0.008 |
| ATF7IP | activating transcription factor 7 interacting protein | 8.135 | 0.026 | 8.179 | 0.021 |
| GPAA1 | glycosylphosphatidylinositol anchor attachment 1 | 8.007 | 0.006 | 8.030 | 0.008 |
| KBTBD11 | kelch repeat and BTB domain containing 11 | 7.922 | 0.015 | 7.877 | 0.018 |
| HINT1 | histidine triad nucleotide binding protein 1 | 7.832 | 0.026 | 7.846 | 0.027 |
| ACO2 | aconitase 2 | 7.799 | 0.013 | 7.817 | 0.016 |
| EIF3G | eukaryotic translation initiation factor 3 subunit G | 7.759 | 0.010 | 7.764 | 0.012 |
| CPNE3 | copine 3 | 7.526 | 0.013 | 7.550 | 0.016 |
| ABL2 | ABL proto-oncogene 2, non-receptor tyrosine | 7.447 | 0.026 | 7.465 | 0.024 |
| ATP6V0E1 | ATPase H+ transporting V0 subunit e1 | 7.407 | 0.034 | 7.431 | 0.040 |
| SCNM1 | sodium channel modifier 1 | 7.364 | 0.046 | 7.398 | 0.045 |
| ARSB | arylsulfatase B | 7.217 | 0.013 | 7.243 | 0.016 |
| ARPC5 | actin related protein 2/3 complex subunit 5 | 7.180 | 0.046 | 7.234 | 0.049 |
| POLR3H | RNA polymerase III subunit H | 7.041 | 0.033 | 7.061 | 0.040 |
| CAPN3 | calpain 3 | 6.914 | 0.037 | 6.911 | 0.052 |
| SF3B6 | splicing factor 3b subunit 6 | 6.869 | 0.030 | 6.929 | 0.021 |
| STRN3 | striatin 3 | 6.729 | 0.047 | 6.755 | 0.060 |
| STAM2 | signal transducing adaptor molecule 2 | 6.554 | 0.047 | 6.573 | 0.060 |

**Supplementary table 5a:** Significantly enriched GO classes in MS nodules vs MS nnNAWM. *Gene ontology analysis was used to find genes of interest based on function. The column count indicates the number of genes likely associated with the GO term and the unadjusted p value is shown*.

| Term | GO class | Count | P |
| --- | --- | --- | --- |
| positive regulation of interferon-gamma production | GO:0032729 | 13 | 1.79e-9 |
| positive regulation of tumor necrosis factor production | GO:0032760 | 13 | 6.65e-8 |
| peptide antigen assembly with MHC class II protein complex | GO:0032755 | 6 | 2.58e-6 |
| positive regulation of interleukin-6 production | GO:0032755 | 11 | 2.93e-6 |
| immunoglobulin production involved in immunoglobulin mediated immune response | GO:0002381 | 6 | 3.61e-6 |
| antigen processing and presentation of exogenous peptide antigen via MHC class II | GO:0019886 | 7 | 4.30e-6 |
| positive regulation of interleukin-8 production | GO:0032757 | 9 | 4.94e-6 |
| antigen processing and presentation of peptide or polysaccharide antigen via MHC class II | GO:0002504 | 6 | 8.71e-6 |
| immune response | GO:0006955 | 22 | 1.43e-5 |
| defense response to virus | GO:0051607 | 14 | 4.11e-5 |
| negative regulation of interferon-gamma production | GO:0032689 | 7 | 4.90e-5 |
| positive regulation of monocyte chemotactic protein-1 production | GO:0071639 | 5 | 7.33e-5 |
| cortical actin cytoskeleton organization | GO:0030866 | 6 | 8.36e-5 |
| erythrocyte differentiation | GO:0030218 | 7 | 1.01e-4 |
| positive regulation of T cell activation | GO:0050870 | 6 | 1.14e-4 |
| innate immune response | GO:0045087 | 22 | 2.22e-4 |
| positive regulation of interleukin-1 beta production | GO:0032731 | 7 | 3.89e-4 |
| positive regulation of T cell proliferation | GO:0042102 | 7 | 3.89e-4 |
| intracellular signal transduction | GO:0035556 | 18 | 3.99e-4 |
| positive regulation of interleukin-12 production | GO:0032735 | 6 | 4.10e-4 |
| inflammatory response | GO:0006954 | 17 | 4.24e-4 |
| positive regulation of NIK/NF-kappaB signaling | GO:1901224 | 7 | 0.001 |
| cellular response to lipopolysaccharide | GO:0071222 | 11 | 0.001 |
| antigen processing and presentation | GO:0019882 | 6 | 0.001 |
| positive regulation of NF-kappaB transcription factor activity | GO:0051092 | 10 | 0.001 |
| receptor internalization | GO:0031623 | 6 | 0.001 |
| cell activation | GO:0001775 | 4 | 0.001 |
| microglial cell activation | GO:0001774 | 5 | 0.001 |
| T cell homeostasis | GO:0043029 | 5 | 0.001 |
| dendritic cell differentiation | GO:0097028 | 4 | 0.001 |
| response to tumor necrosis factor | GO:0034612 | 5 | 0.002 |
| cellular response to diacyl bacterial lipopeptide | GO:0071726 | 3 | 0.002 |
| positive regulation of I-kappaB kinase/NF-kappaB signaling | GO:0043123 | 10 | 0.002 |
| lipopolysaccharide-mediated signaling pathway | GO:0031663 | 5 | 0.003 |
| positive regulation of interleukin-10 production | GO:0032733 | 5 | 0.003 |
| positive regulation of superoxide anion generation | GO:0032930 | 4 | 0.003 |
| phagocytosis | GO:0006909 | 6 | 0.004 |
| negative regulation of T cell proliferation | GO:0042130 | 5 | 0.005 |
| neutrophil activation involved in immune response | GO:0002283 | 3 | 0.006 |
| positive regulation of T-helper 1 cell cytokine production | GO:2000556 | 3 | 0.006 |
| negative regulation of interleukin-10 production | GO:0032693 | 4 | 0.006 |
| defense response | GO:0006952 | 6 | 0.006 |
| positive regulation of phagocytosis | GO:0050766 | 5 | 0.006 |
| positive regulation of interleukin-17 production | GO:0032740 | 4 | 0.006 |
| signal transduction | GO:0007165 | 31 | 0.007 |
| positive regulation of interleukin-4 production | GO:0032753 | 4 | 0.007 |
| extrinsic apoptotic signaling pathway | GO:0097191 | 5 | 0.007 |
| positive regulation of gene expression | GO:0010628 | 16 | 0.010 |
| substrate adhesion-dependent cell spreading | GO:0034446 | 5 | 0.011 |
| positive regulation of transcription from RNA polymerase II promoter | GO:0045944 | 29 | 0.011 |
| calcium-mediated signaling | GO:0019722 | 6 | 0.012 |
| positive regulation of cell migration | GO:0030335 | 10 | 0.014 |
| extrinsic apoptotic signaling pathway in absence of ligand | GO:0097192 | 4 | 0.015 |
| platelet activation | GO:0030168 | 5 | 0.015 |
| organelle transport along microtubule | GO:0072384 | 3 | 0.018 |
| positive regulation of granulocyte macrophage colony-stimulating factor production | GO:0032725 | 3 | 0.020 |
| negative regulation of activated T cell proliferation | GO:0046007 | 3 | 0.020 |
| regulation of tumor necrosis factor-mediated signaling pathway | GO:0010803 | 3 | 0.023 |
| positive regulation of inflammatory response | GO:0050729 | 6 | 0.024 |
| integrin-mediated signaling pathway | GO:0007229 | 6 | 0.024 |
| positive regulation of neuron death | GO:1901216 | 4 | 0.025 |
| apoptotic signaling pathway | GO:0097190 | 5 | 0.026 |
| negative regulation of tumor necrosis factor production | GO:0032720 | 5 | 0.027 |
| positive regulation of protein ubiquitination | GO:0031398 | 5 | 0.028 |
| response to lipopolysaccharide | GO:0032496 | 7 | 0.028 |
| cytokine-mediated signaling pathway | GO:0019221 | 7 | 0.028 |
| late endosome to lysosome transport | GO:1902774 | 3 | 0.028 |
| negative regulation of T cell activation | GO:0050868 | 3 | 0.028 |
| positive regulation of nitric-oxide synthase biosynthetic process | GO:0051770 | 3 | 0.028 |
| positive regulation of B cell proliferation | GO:0030890 | 4 | 0.029 |
| positive regulation of chemokine production | GO:0032722 | 4 | 0.029 |
| animal organ regeneration | GO:0031100 | 4 | 0.029 |
| platelet aggregation | GO:0070527 | 4 | 0.029 |
| regulation of small GTPase mediated signal transduction | GO:0051056 | 6 | 0.029 |
| negative regulation of leukocyte apoptotic process | GO:2000107 | 2 | 0.029 |
| toll-like receptor TLR6:TLR2 signaling pathway | GO:0038124 | 2 | 0.029 |
| methotrexate transport | GO:0051958 | 2 | 0.029 |
| detection of diacyl bacterial lipopeptide | GO:0042496 | 2 | 0.029 |
| regulation of G-protein coupled receptor protein signaling pathway | GO:0008277 | 4 | 0.032 |
| angiogenesis | GO:0001525 | 9 | 0.034 |
| positive regulation of nitric oxide biosynthetic process | GO:0045429 | 4 | 0.036 |
| adaptive immune response | GO:0002250 | 13 | 0.036 |
| B cell differentiation | GO:0030183 | 5 | 0.038 |
| MyD88-dependent toll-like receptor signaling pathway | GO:0002755 | 3 | 0.041 |
| stimulatory C-type lectin receptor signaling pathway | GO:0002223 | 3 | 0.041 |
| MAPK cascade | GO:0000165 | 6 | 0.041 |
| negative regulation of lymphocyte activation | GO:0051250 | 2 | 0.044 |
| reduced folate transmembrane transport | GO:0098838 | 2 | 0.044 |
| nitric oxide metabolic process | GO:0046209 | 2 | 0.044 |
| positive regulation of T cell activation via T cell receptor contact with antigen bound to MHC molecule on antigen presenting cell | GO:2001190 | 2 | 0.044 |
| cellular response to triacyl bacterial lipopeptide | GO:0071727 | 2 | 0.044 |
| positive regulation of glial cell proliferation | GO:0060252 | 3 | 0.045 |
| cell projection assembly | GO:0030031 | 3 | 0.045 |
| transmembrane receptor protein tyrosine kinase signaling pathway | GO:0007169 | 6 | 0.046 |
| positive regulation of protein complex assembly | GO:0031334 | 4 | 0.047 |
| toll-like receptor signaling pathway | GO:0002224 | 3 | 0.048 |

**Supplementary table 5b:** ***Significantly enriched GO classes in MS nodules vs stroke nodules.*** *Gene ontology analysis was used to find genes of interest based on function. The column count indicates the number of genes likely associated with the GO term and the unadjusted p value is shown*.

| Term | GO class | Count | P |
| --- | --- | --- | --- |
| regulation of catalytic activity | GO:0050790 | 13 | 0.001 |
| protein transport | GO:0015031 | 14 | 0.001 |
| Golgi to plasma membrane transport | GO:0006893 | 4 | 0.003 |
| protein targeting to mitochondrion | GO:0006626 | 4 | 0.003 |
| regulation of small GTPase mediated signal transduction | GO:0051056 | 6 | 0.009 |
| membrane fission | GO:0090148 | 4 | 0.009 |
| ER to Golgi vesicle-mediated transport | GO:0006888 | 6 | 0.014 |
| intrinsic apoptotic signaling pathway in response to DNA damage | GO:0008630 | 4 | 0.017 |
| mitochondrial translation | GO:0032543 | 5 | 0.020 |
| regulation of mitochondrial DNA metabolic process | GO:1901858 | 2 | 0.021 |
| actin cytoskeleton reorganization | GO:0031532 | 4 | 0.026 |
| lipid metabolic process | GO:0006629 | 7 | 0.026 |
| Rho protein signal transduction | GO:0007266 | 4 | 0.027 |
| regulation of Rho protein signal transduction | GO:0035023 | 3 | 0.037 |
| response to aluminum ion | GO:0010044 | 2 | 0.043 |
| glial cell apoptotic process | GO:0034349 | 2 | 0.043 |
| T cell homeostasis | GO:0043029 | 3 | 0.047 |
| actin cytoskeleton organization | GO:0030036 | 6 | 0.048 |

**Supplementary table 4:**
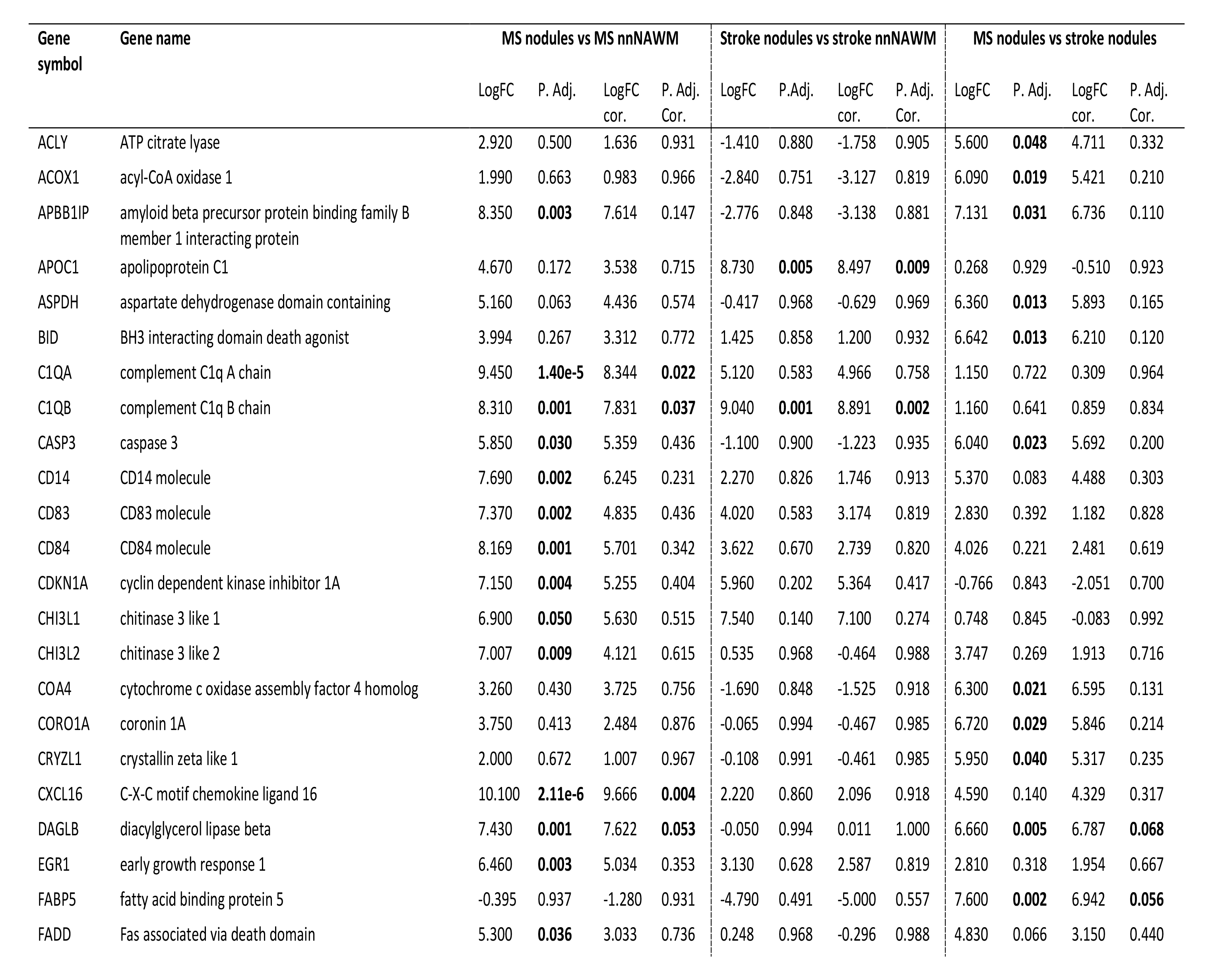
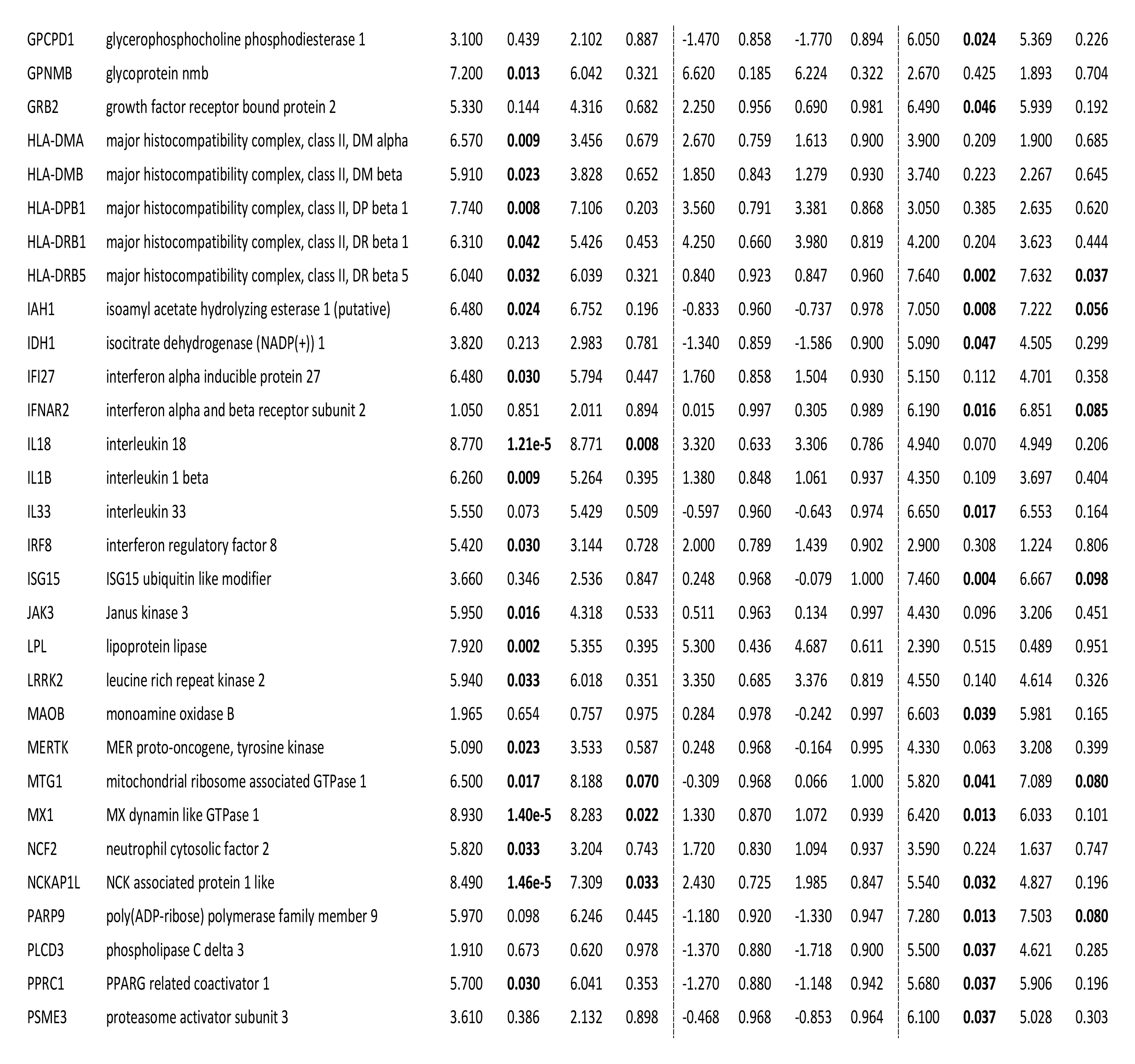
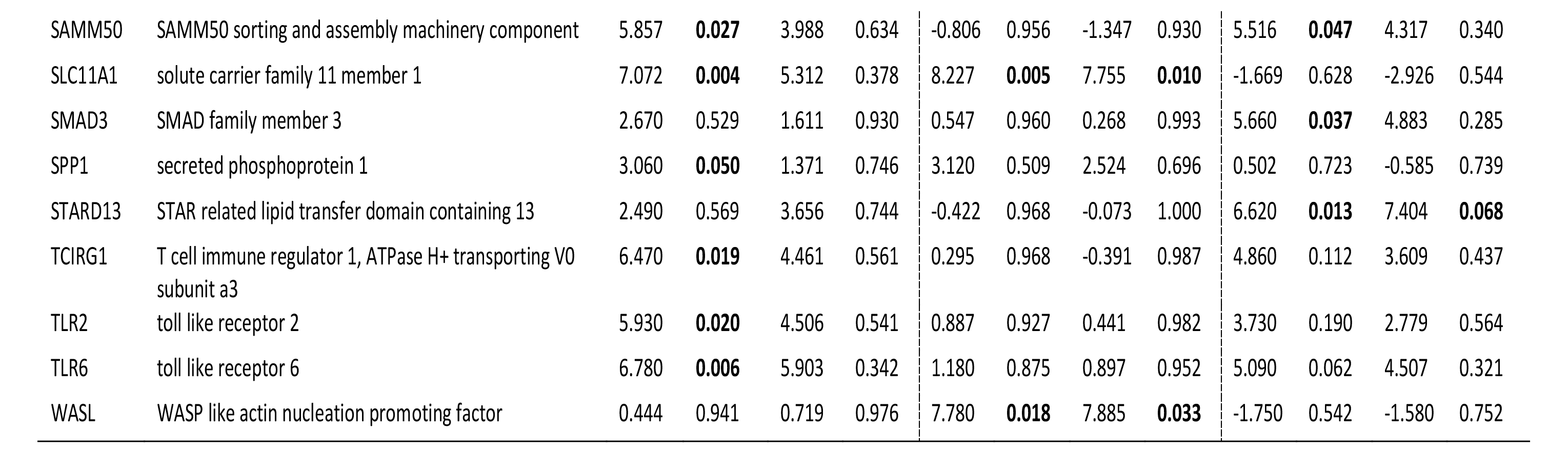
DE genes of interest. *Genes of interest either found through literature or gene ontology analysis are shown for the various comparisons, sorted alphabetically. LogFC and adjusted p values are given without correction for microglia proportion and with correction for microglia proportion. Without correction, p values are considered significant at <0.05. With correction, p values are considered significant at <0.10*.

